# Intestinal tuft cells assemble a cytoskeletal superstructure composed of co-aligned actin bundles and microtubules

**DOI:** 10.1101/2024.03.19.585757

**Authors:** Jennifer B. Silverman, Evan E. Krystofiak, Leah R. Caplan, Ken S. Lau, Matthew J. Tyska

## Abstract

**Background & Aims:** All tissues consist of a distinct set of cell types, which collectively support organ function and homeostasis. Tuft cells are a rare epithelial cell type found in diverse epithelia, where they play important roles in sensing antigens and stimulating downstream immune responses. Exhibiting a unique polarized morphology, tuft cells are defined by an array of giant actin filament bundles that support ∼2 μm of apical membrane protrusion and extend over 7 μm towards the cell’s perinuclear region. Despite their established roles in maintaining intestinal epithelial homeostasis, tuft cells remain understudied due to their rarity (e.g. ∼ 1% in the small intestinal epithelium). Details regarding the ultrastructural organization of the tuft cell cytoskeleton, the molecular components involved in building the array of giant actin bundles, and how these cytoskeletal structures support tuft cell biology remain unclear.

**Methods:** To begin to answer these questions, we used advanced light and electron microscopy to perform quantitative morphometry of the small intestinal tuft cell cytoskeleton.

**Results:** We found that tuft cell core bundles consist of actin filaments that are crosslinked in a parallel “barbed-end out” configuration. These polarized structures are also supported by a unique group of tuft cell enriched actin-binding proteins that are differentially localized along the giant core bundles. Furthermore, we found that tuft cell actin bundles are co-aligned with a highly ordered network of microtubules.

**Conclusions:** Tuft cells assemble a cytoskeletal superstructure that is well positioned to serve as a track for subcellular transport along the apical-basolateral axis and in turn, support the dynamic sensing functions that are critical for intestinal epithelial homeostasis.

**SYNOPSIS:** This research leveraged advanced light and electron microscopy to perform quantitative morphometry of the intestinal tuft cell cytoskeleton. Three-dimensional reconstructions of segmented image data revealed a co-aligned actin-microtubule superstructure that may play a fundamental role in tuft cell function.

## INTRODUCTION

Epithelial tissues are composed of a diverse array of cell types including transporting, sensory, and secretory cells, which work together to support physiological homeostasis. However, not all cell types are equally represented within the epithelium, and certain populations are extremely rare. One such rare cell type is the tuft cell (e.g. less than 1% of cells in the small intestine), which is broadly characterized as a chemosensory cell and found in the thymus, gastrointestinal, respiratory, and urogenital tracts [1–5]. Perhaps because of their rarity, our understanding of their basic biological features and how they support epithelial tissue physiology remain understudied.

Tuft cells from diverse tissues perform similar functions dependent on G-protein coupled receptors (GPCRs) sensing ligands specific to their biological niche. In the small intestine, tuft cells sense bacteria, protists, and helminths using a variety of GPCRs including succinate receptor 1 and free fatty acid receptor 3 [6, 7]. Through the canonical taste receptor pathway, transient receptor potential cation channel subfamily M member 5 (TRPM5) causes an influx of sodium that depolarizes the cell [8]. Downstream, in pathways that are not yet defined, depolarization causes the secretion of tuft cell effectors including interleukin 25 (IL-25), cysteinyl leukotrienes (CysLTs), prostaglandin D2 (PGD2), and acetylcholine (Ach) [9–12]. These effectors trigger a type 2 innate lymphoid cell response which, in the intestine, results in tuft cell hyperplasia and aids in the subsequent clearance of parasites and barrier maintenance [9–12]. Interestingly, tuft cells may be heterogenous within individual tissues, with evidence for two broad subtypes. Type-1 tuft cells are classified as neuron-like and express choline acetyltransferase, an enzyme required for synthesis of acetylcholine, whereas type-2 tuft cells are classified as immune-like and express immune-related genes such as pan-immune marker CD45/Ptprc [13]. Although animal studies are beginning to reveal how tuft cells contribute to epithelial physiology, the subcellular mechanisms underlying tuft cell function in these diverse contexts remains poorly understood.

Tuft cells were first characterized by their morphology as ‘peculiar cells’ with a small apical surface that presents a large ‘tuft’ of protrusions [14, 15]. Superficially, tuft cell protrusions exhibit a fingerlike morphology resembling microvilli. However, early electron microscopy (EM) data revealed some unique features, e.g. tuft protrusions are supported by ‘giant’ actin-rich structures, which protrude ∼2 µm from the cell surface wrapped in apical membrane and extend many microns deep into the cytoplasm, down to the perinuclear region [2, 16, 17]. These imaging studies also noted microtubules beneath the tuft cell apical surface [18, 19], which is common for polarized epithelial cell types. Interestingly, EM volume reconstructions of single tuft cells revealed that the sub-apical cytoplasm contains a large ‘tubulovesicular structure’, although any connection between this structure and the cytoskeletal features alluded to above remains unclear [20]. Later studies using light microscopy and single cell RNA sequencing (scRNAseq), reported enrichment of specific cytoskeletal components in tuft cells, including acetylated tubulin [4], actin bundling protein fimbrin [2], actin and Akt binding protein girdin [21], and actin binding protein advillin [22–24]. Advillin is part of a well characterized family of actin bundlers and has high homology (59%) with villin, a major bundler in enterocyte microvilli [25].

Although previous studies hinted at the unique organization and composition of the tuft cytoskeleton, important questions remain unanswered. For example, the molecular mechanisms that drive the formation of giant actin bundles or promote the packing of apical protrusions into a tuft remain undefined. How this cytoskeletal specialization supports the physiological functions of tuft cells also remains unclear. Answers to these questions will be essential for understanding how tuft cells are adapted to support homeostasis through sensing and secretion. To address these gaps, we leveraged quantitative light and electron microscopy, in combination with trainable image segmentation, with the goal of building detailed three-dimensional views of the cytoskeletal structures underlying the tuft, in tuft cells in the small intestine. To work around the problem of tuft cell rarity in this tissue, we took advantage of succinate treatment to increase tuft cell specification [26]. Using these approaches, we defined the organization and packing of actin filaments within tuft cell protrusions, and characterized the subcellular distribution of cytoskeletal proteins that exhibit tuft cell specific enrichment. Our studies identified a striking array of microtubules that exhibit interdigitating co-alignment with giant actin bundles, forming a superstructure of parallel cytoskeletal polymers that extend from the apical surface down to the perinuclear region. Extensive decoration of this superstructure with multi-vesicular bodies and other small membranous organelles suggests that this unique architecture might support vectorial transport along the apical-basolateral axis. These discoveries advance our knowledge of fundamental tuft cell biology and suggest models for how the unique cytoskeleton found within this rare cell type might contribute to sensing and secretion, subcellular activities that are essential for intestinal homeostasis.

## RESULTS

### The tuft cell ‘tuft’ contains ∼100 giant actin bundles

Our review of the tuft cell literature revealed a lack of cohesive terminology for describing the sub-cellular features of this unique cell type. Herein, we propose and apply a simple nomenclature when referring to specific structures within the tuft, defined here as the full array of surface protrusions and their supporting core actin bundles (Figure 1A). Our hope is that these terms will be adopted by others in the field to facilitate clear and coherent communication. To begin defining the structural architecture of the tuft, we used confocal and super-resolution light microscopy to visualize the tuft cytoskeleton. As a model for these studies, we took advantage of mouse small intestinal tissue sections and whole mount preparations; these allowed us to visualize single tuft cells in both lateral and *en face* (top down) views. We first performed high-resolution confocal imaging of tissue sections labeled with fluorescent phalloidin to visualize F-actin. Lateral views of single tuft cells revealed arrays of long, continuous actin bundles ranging from 5 to 12 µm in length (median: 7) (Figure 1B,C), up to ∼10 times longer than the core bundles that support microvilli on neighboring enterocytes [27]. Close inspection of the phalloidin signal along these giant actin bundles revealed that they extend as a single, continuous structure from the distal end of the supported protrusion, down to the perinuclear region (Figure 1B). Imaging whole mount sections captured just under the apical surface enabled us to define the number of actin bundles per cell (Figure 1D). Tuft cells contained a median of 101 core bundles (Figure 1E) and linescans drawn along the bundle axis show that phalloidin signal tapers toward the rootlet end (Figure 1F). Although phalloidin intensity is generally proportional to actin filament density, whether such loss of signal towards the rootlet end represents a decrease in filament number, or alternatively, inhibition of phalloidin binding by enrichment of an actin-associated protein remains unclear.

**Figure 1.**
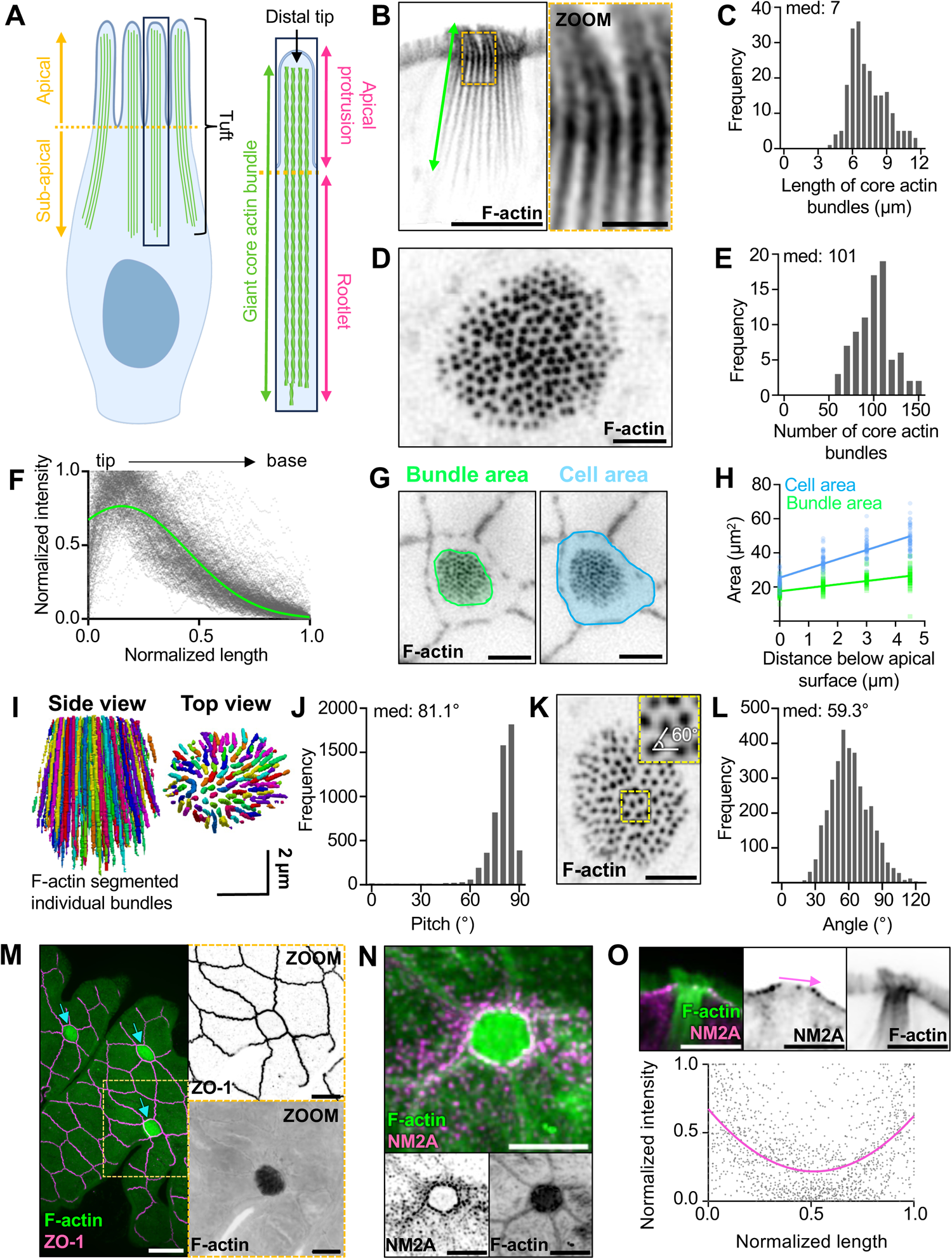
The tuft cell ‘tuft’ contains over a hundred of giant actin bundles. **A)** Cartoon depicting nomenclature for giant core actin bundles in tuft cells. **B)** Inverted single channel, single Z-slice, Airyscan image of lateral tissue section of tuft cell. Actin marked with phalloidin (scalebar = 5 µm, zoom = 1 µm). **C)** Frequency diagram of core bundle length as depicted in Fig 1B (n = 27 tuft cells over 3 mice). **D)** Max intensity projection (MaxIP) spinning disk confocal (SDC) image using *en face* whole-mount tissue of a tuft cell captured beneath the apical membrane with phalloidin staining (scalebar = 2 µm). **E)** Frequency diagram of the number of core actin bundles in tuft cells (n = 81 tuft cells over 3 mice). **F)** Using lateral sections of frozen tissue, linescans of phalloidin intensity were drawn from tip to base of core actin bundles. Raw linescan data shown in gray with a curve fit to data shown in yellow. (n = 218 bundles over 3 mice). **G)** Single z-slice SDC image of *en face* whole mount tissue (scale bar = 5 µm). **H)** Simple linear regression measuring cell or bundle area (as shown in Fig 1G) at 1.5 µm, 3 µm, and 4.5 µm beneath the apical surface. Slope bundle area = 2.013. Slope cell area = 5.412 (n = 54 tuft cells over 3 mice). **I)** 3D projection of giant core actin bundles in a tuft cell using Trainable WEKA segmentation. **J)** Pitch of individual actin bundles relative to the long axis of the cell (90° = vertical orientation) using WEKA segmented data from Fig 1I (n = 35 tuft cells over 3 mice). **K)** MaxIP SDC image of *en face* whole-mount tissue section with showing sub-apical giant core bundles (scalebar = 2 µm). **L)** Frequency diagram of angle measurements between neighboring bundles as shown in zoom inset Fig 1K (n = 3453 measurements made in 25 tuft cells over 3 mice). **M)** MaxIP SDC image of *en face* whole mount tissue with ZO-1 (magenta) and actin marked by phalloidin (green) (scale bar = 5 µm). **N)** MaxIP SDC image of *en face* frozen tissue section with NM2A (magenta) and actin marked with phalloidin (green) (scalebar = 5 µm). O) Above: single Z-slice image of lateral frozen tissue section with NM2A (magenta) and actin marked with phalloidin (green) (scalebar = 5 µm). Below: linescans measuring the intensity of NM2A across the apical surface of tuft cells as shown by the pink arrow in the image above (n = 33 tuft cells over 3 mice)

Within the small intestine, tuft cells exhibit a fusiform (i.e. bottle shaped) morphology, with a narrow apical surface and wider cell body [17, 28]. To determine whether the array of core actin bundles is shaped by the overall morphology of the cell, we measured the cross-sectional area of the cell body relative to the area occupied by bundles, moving down from the apical surface in 1.5 µm increments (Figure 1G). Right at the apical surface, bundle area and cell area are similar (Figure 1H), suggesting that the junctional actomyosin belt constrains bundle spacing in this plane. However, both bundle area and cell area progressively increased below the apical surface, although the rate of increase was higher for the cell body, indicating that the width of cell body does not limit bundle spreading deeper in the cell (Figure 1H). To better visualize the orientation of individual bundles relative to the apicobasal axis, bundles were segmented via trainable Waikato Environment for Knowledge Analysis (WEKA) segmentation [29], thresholds were applied to the generated probability maps to reduce noise (see Methods), and structures were then rendered in 3D (Figure 1I). Bundles in these reconstructions demonstrated a median tilt of 81.1°, splaying slightly outward from the apicobasal axis (90°) (Figure 1J), again with narrowest point of constriction right at the apical surface. We also examined the packing organization of bundles within the tuft (Figure 1K). Enterocyte microvilli exhibit hexagonal packing, which maximizes the number of protrusions and thus, apical membrane holding capacity per cell. In the tuft, measurements of the angle formed by groups of three adjacent protrusions revealed a median of 59.3° suggestive of hexagonal packing, although the spread around this value was large, indicating areas of fluid packing (Figure 1L). To gain insight on mechanisms that constrict giant actin bundle packing at the apical surface, we examined the actomyosin belt that forms at the level of junctional complexes [30]. Tuft cells have long been identified by their small apical profiles (*en face* area) [16, 17], and our measurements of *en face* cell surface area using tight junction marker, zonula occludens 1 (ZO-1) (Figure 1M), did reveal that tuft cells exhibit significantly smaller and more circular apical areas than neighboring enterocytes (Figure S1A, B). The small radius of the tuft cell apical profile is suggestive of elevated contractility in the junctional actomyosin belt [31]. Intestinal epithelial cells express three distinct non-muscle myosin-2 isoforms - NM2A, NM2B, and NM2C [32], with NM2A supporting the highest levels of contractility [33]. In the case of tuft cells, *en face* images showed NM2A enrichment in the junctional actomyosin belt (Figure 1N) and lateral views of tissue sections also revealed puncta enriched at the level of junctional contacts (Figure 1O); a similar distribution was observed for NM2C (Figure S1C, D). Relative to enterocytes, tuft cells express similar levels of NM2A (Figure S1E, F) but much lower levels of NM2C (Figure S1G, H). These data suggest that the constrained diameter of the tuft cell apical surface and the tuft are primarily due to a high NM2A/NM2C ratio.

### Giant actin bundles contain ∼100 hexagonally packed actin filaments

We next sought to define the organization of actin filaments within individual giant core actin bundles. We used transmission EM (TEM) to collect *en face* cross-sectional views of the tuft, which enabled us to resolve individual actin filaments in core bundles. To overcome the difficulty of capturing rare tuft cells in ultrathin sections, we increased the number of tuft cells using an established model of succinate-driven tuft cell hyperplasia [34]. With this approach, we found that bundles (Figure 2A) contain ∼100 filaments (median: 101.5) (Figure 2B), in contrast to enterocyte microvilli, which contain only 20-30 [27]. We also noted that the distance from the core actin bundle to the surrounding membrane was spaced at a median of 16.8 nm (Figure 2C), which is consistent with the dimensions of known membrane-actin linkers, including class 1 myosins [35].

**Figure 2.**
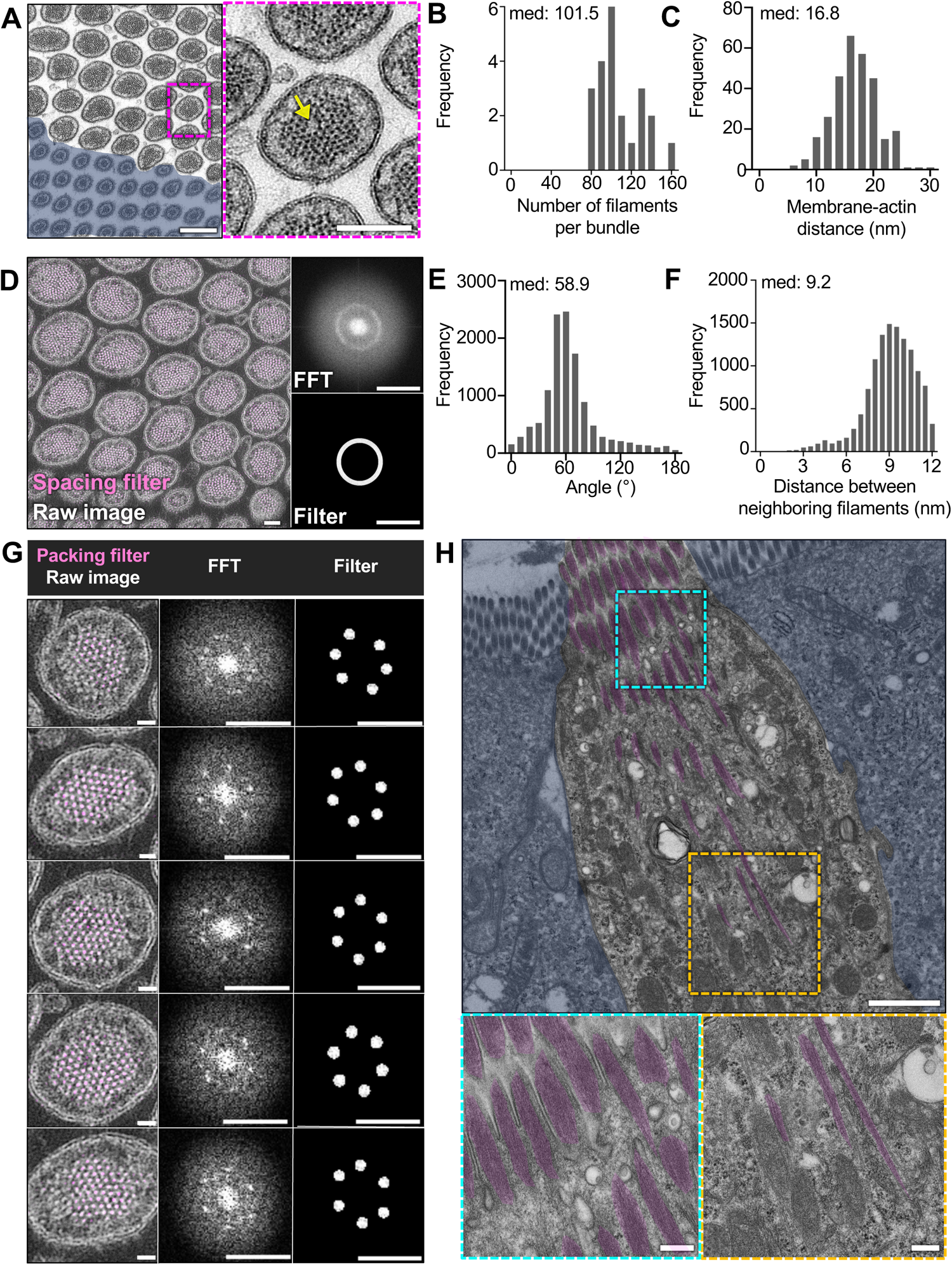
Giant actin bundles contain ∼100 hexagonally packed actin filaments. **A)** Transmission electron microscopy (TEM) of ultrathin tissue section depicting tuft cell apical protrusions imaged *en face* (scalebar = 200 nm) and enterocyte masked in blue, zoom inset on right (scalebar = 100 nm). Yellow arrow points to instance of actin dislocation. **B)** Frequency diagram of number of filaments using ultrathin sections such as Fig 2A (n = 22 bundles over 3 tuft cells). **C)** Frequency diagram distance between outer protrusion membrane to actin measured at 5 separate places in each bundle (n = 3 tuft cells). **D)** TEM of tissue section tuft cell apical protrusions imaged *en face*. Map of evenly spaced actin filaments (magenta), Fast Fourier transform (FFT) (top right), bandpass filter (bottom right) (scalebar = 100 nm). **E)** Frequency diagram of the angle between nearest neighbor filaments within a 12 nm radius (n= 22 bundles over 3 tuft cells). **F)** Frequency diagram of the distance between neighboring filaments identified in E (n = 22 bundles over 3 tuft cells). **G)** Map of hexagonally packed filaments on 5 different protrusions taken from Fig 2D. FFT (top right), bandpass filter (bottom right) (scalebar raw image = 20 nm, FFT and filter scalebar = 20 nm^−1^). **H)** TEM of tissue section depicting mostly lateral section of a tuft cell with enterocytes masked in blue. Giant core bundles highlighted in green (scalebar = 1 µm). Zoom insets on bottom show the base of the rootlets (left) and core bundles near the apical surface (right) (scalebars = 200 nm).

To further examine the arrangement of giant core bundle actin filaments, we used Fourier analysis to create frequency space maps that highlight dominant spacing features. In image fields containing many bundle cross-sections, Fourier maps were indicative of uniform filament-filament spacing (Figure 2D). Fourier patterns were also used to create a filter that was then overlayed with the original image (magenta), to highlight all filaments with uniform spacing (Figure 2D). Virtually all filaments were highlighted showing uniform spacing. Because such uniform inter-filament spacing is typically associated with hexagonal packing patterns, we performed a similar analysis on individual bundles to examine this possibility. Here, we generated Fourier filters to highlight only filaments with hexagonal packing (magenta) (Figure 2G). In all bundles analyzed, the majority of actin filaments were packed hexagonally. We then used Trackmate, a FIJI plugin, to identify filament center coordinates with sub-pixel precision (Figure S2A). Counting nearest neighbors within a 12 nm radius (approximately twice the width of a single filament), it was evident that hexagonally packed filaments, indicated by 6 nearest neighbors, were enriched in the center of the bundle. Filaments at the edge of the bundle deviated from perfect hexagonal packing, which reduced the number of nearest neighbors to 5 (Figure S2B). Additionally, the angle formed by groups of three neighboring filaments was close to 60° (median: 58.9°) (Figure 2E), which is consistent with hexagonal packing. This analysis also revealed that the mean center-to-center distance between filaments was ∼9 nm (median: 9.2 nm) (Figure 2F). Finally, we noted that some bundles contained packing anomalies (dislocations) where filaments were entirely missing (yellow arrow in Figure 2A), although this did not appear to disrupt the organization of filaments around these voids. To visualize the filaments within giant actin bundles along their length, we examined lateral tissue sections via TEM (Figure 2H). Using this approach, it was difficult to capture complete views of entire bundles. However, we did observe that bundle thickness and the number of filaments decrease toward the rootlet ends, which is consistent with the decay in phalloidin signal (Figure 1F). The filament packing patterns revealed in these ultrastructural studies hold implications for understanding mechanisms that drive tuft formation, as we discuss in more detail below.

### Filaments in giant core actin bundles exhibit uniform ‘barbed-end out’ polarity

Based on our TEM images, core bundle actin filaments appear to be crosslinked tightly, in parallel. However, the orientation of individual filaments within these bundles has not been defined. Filament orientation in this scenario is important because it will constrain models for how these structures contribute to subcellular function. Bundles consisting of filaments with uniform orientation are well-suited for supporting directional motor-driven transport [36], whereas mixed orientation bundles are typically associated with contractile functions [37]. To determine bundle orientation, we stained mouse small intestine with epidermal growth factor receptor pathway substrate 8 (EPS8), an actin binding protein that marks barbed-ends of filaments in bundles with uniform orientation [38]. Previous studies established that this factor is highly enriched at the distal tips of microvilli, filopodia, and stereocilia, where the barbed-ends of uniformly oriented filaments reside [39–41]. Confocal images revealed that tuft cells do exhibit some distal tip targeting of EPS8, although linescan analysis showed significantly lower levels than enterocytes (Figure 3A, C). From this perspective, we then stained for other EPS8-like molecules including EPS8-like 2 (EPS8L2), which has a domain organization similar to EPS8 [42]. Relative to neighboring enterocytes, EPS8L2 is highly enriched in tuft cells where it localizes specifically to the distal tips of apical protrusions (Figure 3B, C). Analysis of single cell RNAseq data also confirmed the higher level of enrichment for EPS8L2 vs. EPS8 in tuft cells (Figure S3A, B). Although there is no direct biochemical evidence confirming EPS8L2 as a barbed-end binder, this factor has been shown to target the tips of row 2 stereocilia in hair cells of the inner ear [43], which is consistent with barbed-end binding. We also expressed an EGFP-tagged version of EPS8L2 in HeLa cells and observed strong localization to the tips of filopodia (Figure 3D), providing additional support for barbed-end binding. Taken together, these data suggest that the tuft cell giant actin bundles are composed of filaments organized with uniform ‘barbed-end out’ polarity.

**Figure 3.**
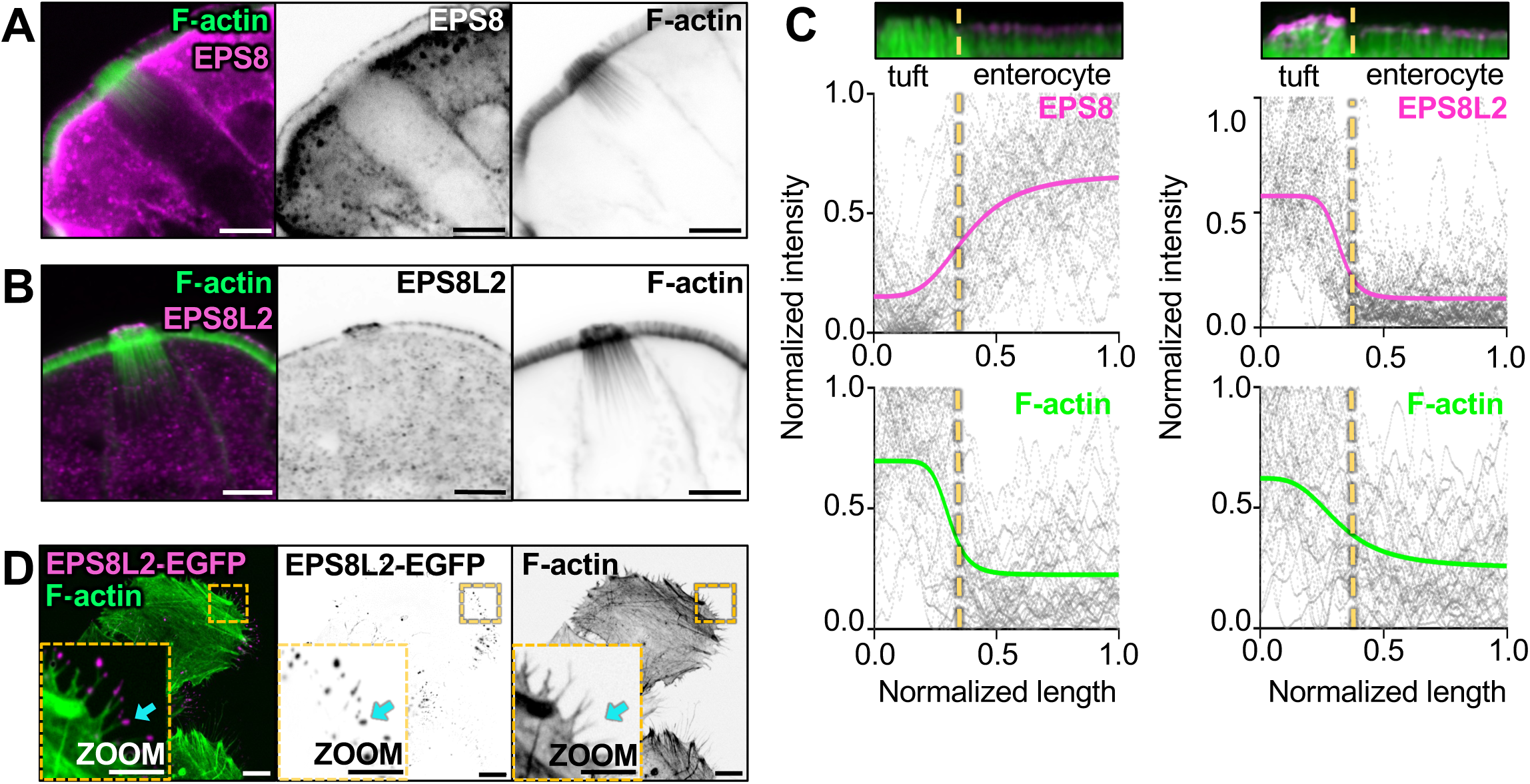
Filaments in giant actin bundles exhibit uniform ‘barbed end out’ polarity. **A)** MaxIP of laser scanning confocal image of EPS8 in tuft cells. Actin marked with phalloidin (scalebar = 5 µm). **B)** MaxIP of laser scanning confocal image of EPS8L2 in tuft cells. Actin marked with phalloidin (scalebar = 5µm). **C)** Linescans drawn over a MaxIP image of the apical tuft and neighboring enterocyte microvilli, measuring EPS8 or EPS8L2 intensity in tuft cells and enterocytes, with accompanying phalloidin intensity. Images provided below as reference. Raw data for all graphs shown in gray, line fit (sigmoidal 4PL) in color. The line fit for EPS8 indicates a bottom plateau of 0.15 normalized intensity in tuft cells and a top plateau of 0.66 normalized intensity in enterocytes. The line fit for EPS8L2 indicates a bottom plateau of 0.13 normalized intensity in enterocytes and a top plateau of 0.57 normalized intensity in tuft cells. (n = EPS8: 37 tuft cells over 3 mice; EPS8L2: 33 tuft cells over 3 mice). **D)** MaxIP SDC image of HeLa cells expressing EPS8L2-EGFP, cyan arrow points to filopodia tip (scale bar = 2 µm).

### Giant core actin bundles show regionalization of tuft cell enriched actin-binding proteins

Cells form higher-order actin structures using a vast array of actin-binding proteins, some of which cross-link and stabilize filaments. For example, stereocilia core bundles contain espin-1, fimbrin/plastin-1, and fascin-2 [44–46] whereas enterocyte microvilli contain espin, villin, fimbrin, and mitotic spindle positioning protein (MISP) [47–50]. Recent research also indicated that these bundlers occupy distinct ‘neighborhoods’ relative to the ends of the bundle [50, 51]. In tuft cells, previous studies identified specific expression of actin-binding proteins including fimbrin, advillin, and girdin (Y1798) [2, 21, 23, 24]. By screening scRNAseq datasets, we also identified additional tuft cell enriched filament bundling candidates including espin and LIM domain and actin binding 1 (LIMA1) (Figure S3C, D). To better understand the localization of espin and LIMA1 within tuft cells, we used super-resolution Zeiss Airyscan microscopy to image immunostained tissue sections. For spatial points of reference, samples were also stained with phalloidin to highlight all F-actin in the tuft, and advillin, an established tuft cell marker [23, 24]. Linescans were drawn along individual core bundles (from distal tip to rootlet) and the signal intensity along that axis was plotted from multiple tuft cells. Advillin displayed high level enrichment in tuft cells, with strong signal at the protruding ends of giant actin bundles that decayed toward the rootlets (Figure 4A, B). Consistent with scRNAseq data, espin was also strongly localized to the protruding ends of giant actin bundles (Figure 4C, D & S3C). Interestingly, LIMA1 was uniquely restricted to the rootlets and maintained a high intensity down to the most proximal ends (Figure 4E, F). Thus, espin and LIMA1 are newly identified components of tuft cell giant actin bundles, and the regionalization of these factors may hold importance for understanding the function and/or mechanical properties of the rootlet vs. protruding bundle segments.

**Figure 4.**
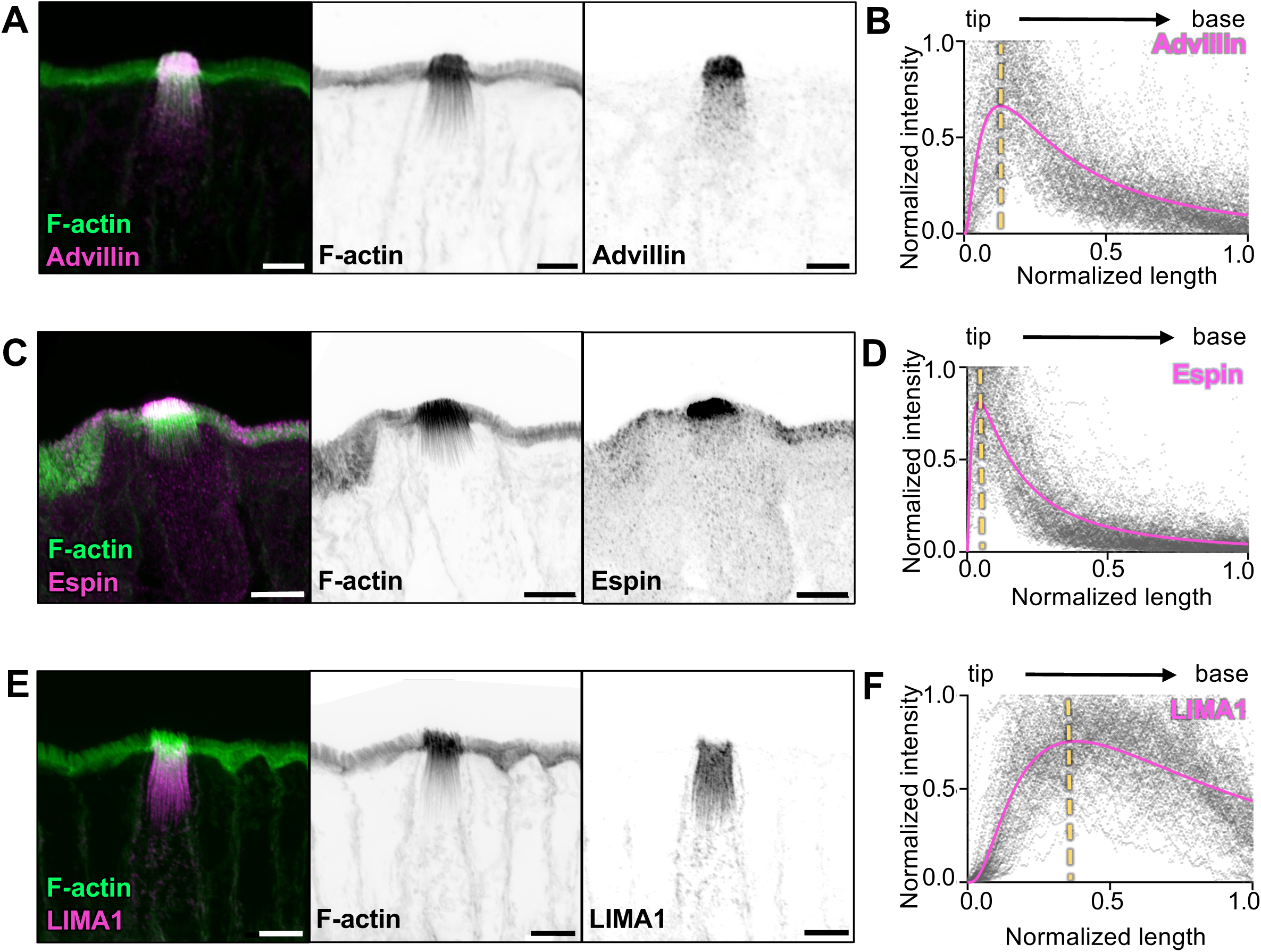
Giant actin bundles show regionalization of tuft cell enriched actin-bundling proteins. **A, C & E)** MaxIP Airyscan image of lateral frozen tissue section and immunostained for actin-binding proteins, (A) advillin, (B) espin, and (C) LIMA1 in addition to actin marked with phalloidin (scalebar = 5 µm). **B, D & F)** Graph of linescans drawn from MaxIP images from apical tip to base of core bundle, measuring intensity of actin binding proteins. Raw values depicted in gray, line fit (lognormal) in magenta, yellow line indicates peak of line fits (n = espin, 33 tuft cells; advillin, 29 tuft cells; LIMA1, 32 tuft cells, over 3 mice).

The cylindrical morphology of surface protrusions such as microvilli is maintained by high levels of linker proteins, including class 1 myosins and ezrin, which stabilize membrane-actin interactions [52, 53]. From this perspective, we sought to determine if tuft cells shared similar membrane-actin linking proteins. Interestingly, scRNAseq data showed a strong enrichment of myosin-1b in tuft cells compared to enterocytes (Figure S3E). As one of eight class 1 myosins expressed in mice, myosin-1b is a tension sensitive motor implicated in vesicle secretion [54]. Immunostaining revealed that myosin-1b is strongly expressed in tuft cells, where it localizes to the protruding ends of giant actin bundles (Figure 5A, B). Ezrin was also detected in tuft cells (Figure S3F) and its expression was similarly restricted to the apical tuft (Figure 5C, D). Ezrin is autoinhibited, but it can be activated through the activity of SLK/LOK kinases, which phosphorylate residue Y565 (pEzrin) [55]. Interestingly, whereas total ezrin levels were similar between tuft cells and enterocytes, tuft cells had significantly higher levels of pEzrin in apical protrusions (Figure 5E-G) suggesting that higher levels of membrane-cytoskeleton adhesion may be needed to achieve the larger dimensions of protrusions in the tuft. Collectively, these immunostaining results highlight a unique complement of actin associated factors that likely cooperate to drive giant bundle assembly and organization within the tuft.

**Figure 5.**
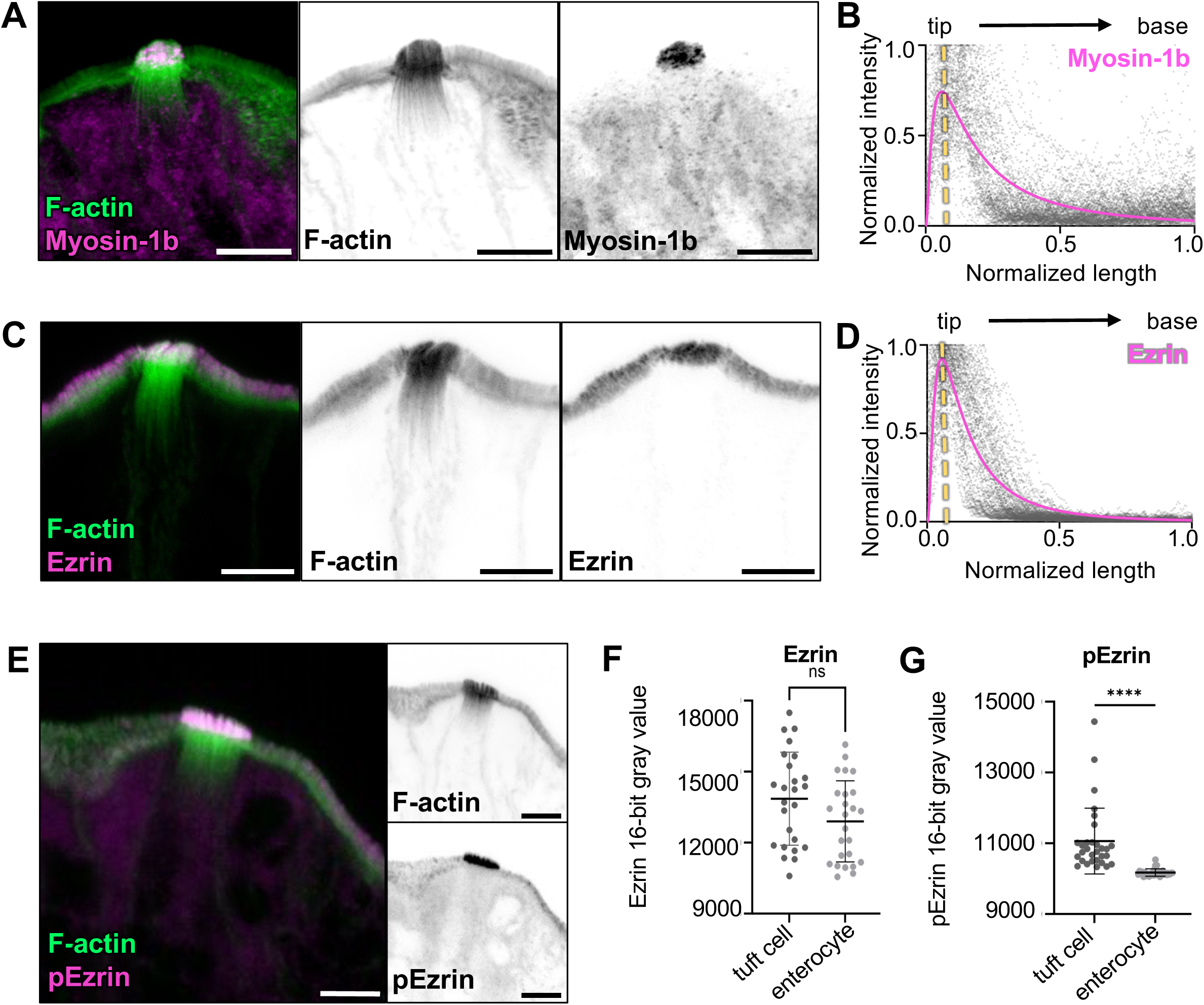
Giant actin bundles show regionalization of tuft cell enriched membrane-actin linking proteins. **A, C, & E)** MaxIP Airyscan image of lateral frozen tissue section and immunostained for actin-binding proteins and actin marked with phalloidin (scalebar = 5 µm). **B & D)** Graph of linescans drawn from MaxIP images from apical tip to base of core bundle, measuring intensity of actin binding proteins. Raw values depicted in gray, line fit (lognormal) in magenta, yellow line indicates peak of line fits (n = myosin 1b, 30 tuft cells; ezrin, 27 tuft cells, over 3 mice). **F)** Mean ezrin intensity in tuft cells versus neighboring enterocytes, using a sum intensity projection, unpaired t-test, p = 0.0723, Error bars denote mean ± SD, (n = 25 tuft cells over 3 mice). **G)** Mean pEzrin intensity in tuft cells versus neighboring enterocytes, unpaired t-test, p<0.001 (n = 30 tuft cells over 3 mice). Error bars denote mean ± SD.

### Giant actin bundles co-align and interdigitate with acetylated microtubules

Previous transcriptomic analysis confirmed an enrichment of tubulin in tuft cells [22] and TEM imaging of ultrathin sections also noted the vertical orientation of microtubules between core actin bundles [16, 18]. Confocal microscopy using markers for tubulin or acetylated tubulin (Ac-tubulin), a post-translationally modified version of tubulin associated with long-lived microtubules [56], also confirmed the presence of long microtubule tracks in tuft cells [4, 19, 57]. Yet how microtubules in the tuft are arranged relative to giant core actin bundles remains unclear. To address this, we stained for Ac-tubulin in frozen tissue sections and imaged these samples using confocal microscopy. Consistent with previous work, we observed that microtubules enriched in Ac-tubulin were oriented along the vertical axis of tuft cell but excluded from apical protrusions (Figure 6A). This microtubule network also extended several microns past the rootlet ends of giant actin bundles, although the overall length of both cytoskeletal networks was comparable (Figure 6B). Like actin filaments, microtubules are polarized polymers, and in polarized epithelial cells (e.g. enterocytes), they generally extend their minus ends towards the apical surface [58, 59]. However, when we stained for dynein heavy chain to mark microtubule minus ends, we unexpectedly found that tuft cells exhibit minimal dynein enrichment at the apical surface relative to neighboring enterocytes (Figure 6C, D), although we noted some dynein labeling towards the perinuclear compartment and strong labeling in the basolateral region (Figure 6E). These data suggest that tuft cells build stable microtubule arrays that extend their plus ends toward the apical surface.

**Figure 6.**
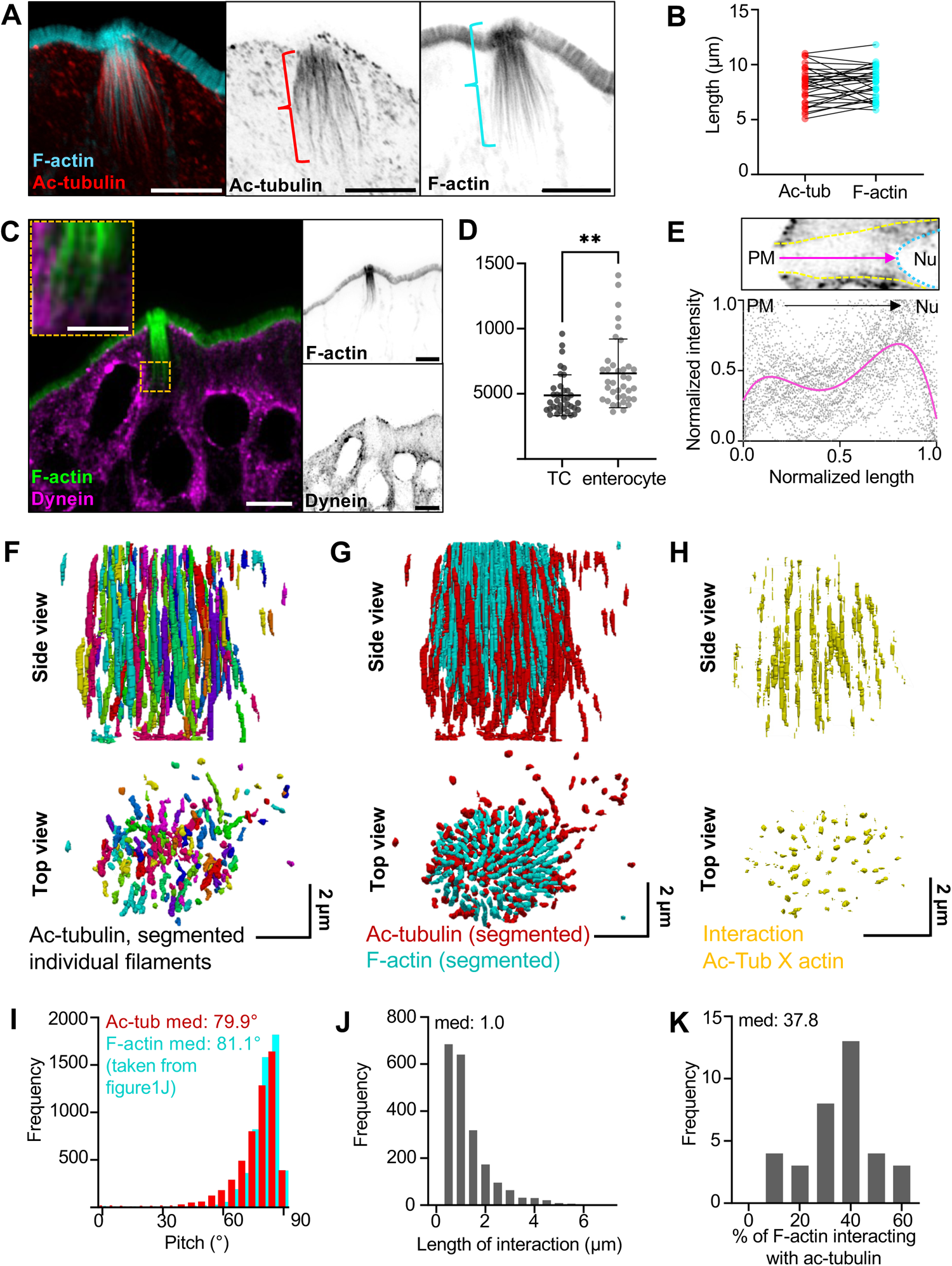
Giant actin bundles co-align and interdigitate with acetylated microtubules. **A)** MaxIP Airyscan image of lateral tissue section (scalebar = 5 µm). **B)** Graph of length of microtubule network versus actin network in tuft cells, measurements demonstrated in Fig 7A. Paired T-test, p = 0.3472. **C)** MaxIP SDC image of lateral tissue section with dynein immunostaining. Actin marked with phalloidin (scalebar = 5 µm, zoom scalebar = 2 µm). **D)** Mean dynein intensity measurements taken from sum intensity projections at the apical surface of tuft cells versus neighboring enterocytes, unpaired t-test, p = 0.0013, Error bars denote mean ± SD, (n = 37 tuft cells over 3 mice) **E)** Graph of linescan showing dynein intensity taken from MaxIP images at the apical surface of tuft cell to top of nucleus. Image above depicts route of linescan (PM = plasma membrane, Nu. = nucleus). Raw data shown in gray, and line fit (fourth-order polynomial) in magenta (n = 38 tuft cells over 3 mice). **F)** 3D projection of acetylated tubulin network in a tuft cell using Trainable WEKA segmentation. **G)** 3D projection of both actin and acetylated microtubule networks, taken from trainable WEKA segmented data. **H)** Interaction between actin and microtubules considered as areas of overlap or immediate adjacency between both cytoskeletal networks and created via dilation of core actin bundles (x4) using the segmented data in Fig 6G. **I)** Frequency diagram showing pitch of microtubules using segmentation from Fig 6F, data on actin pitch taken from Figure 1J (n = 35 tuft cells over 3 mice). **J)** Frequency diagram of the length of individual interactions from Figure 6H (n = 35 tuft cells over 3 mice). **K)** Frequency diagram of the percentage of total actin within a tuft cell that is interacting with microtubules. Calculated as total length of interaction (Figure 6J) divided by total length of actin per tuft cell (not shown).

Similar to our analysis of giant core actin bundles, we used trainable WEKA segmentation to generate a 3D reconstruction of the Ac-tubulin signal in tuft cells (Figure 6F). The overall positioning, organization, and tilt of the microtubules (relative to the apicobasal axis) was remarkably similar to that of giant core actin bundles (microtubule median: 79.9°; F-actin median: 81.1°) (Figure 6I), suggesting the possibility of physical interactions between these two cytoskeletal networks. The strong co-alignment of giant actin bundles and Ac-tubulin signal is also apparent when superimposing their reconstructions (Figure 6G). To better visualize the alignment of these networks, we generated a separate map showing actin bundle segments that were overlapping or immediately adjacent to microtubules (see interaction map, Figure 6H). It is important to note that our 3D maps were drawn with conservative thresholding parameters (see Methods), so the full scale of microtubule-actin overlap or contact may be underestimated. Despite this, our analysis revealed extended segments of contact ranging up to 4 μm in length (median: 1.0 µm) (Figure 6J). Moreover, within each tuft cell, a large percentage (median: 37.8%) of the core actin bundles are marked as interacting with the microtubules using this approach (Figure 6K). Taken together, these findings reveal that tuft cell giant actin bundles co-align and interdigitate with an array of stable microtubules, forming a cytoskeletal superstructure that extends from the apical surface, down through the sub-apical cytoplasm, to the perinuclear region.

### Membranous organelles associate with cytoskeletal polymers in the tuft

The highly ordered, interdigitated network of acetylated microtubules and giant core actin bundles within the sub-apical cytoplasm is uniquely positioned and organized to support motor-driven trafficking of cargoes between the perinuclear region and apical tuft. Consistent with this proposal, TEM revealed numerous small vesicles and membranous organelles in the submicron range tightly associated with giant actin bundles and co-aligned microtubules (Figure 7A). These include both electron lucent vesicles (green arrow) as well as potential multivesicular bodies (MVBs, blue arrow). We also observed vesicles that appeared compressed by the surrounding core bundles, implying direct physical association (see green arrow). Interestingly, TEM images showing cross-sections through apical protrusions revealed an abundance of extracellular vesicles (EVs) situated between apical protrusions. These EVs were similar in dimensions and appearance to the smaller vesicles noted in the sub-apical cytoplasm (Figure 7B, cyan), suggesting that they might be released by tuft cells into the extracellular space. To further examine the possibility that cytoskeletal polymers in the tuft support trafficking of organelles to or from the apical surface, we used light microscopy to examine the distribution of potential membrane protein cargoes. Here we took advantage of mice expressing endogenously tagged cadherin related family member 5 (CDHR5), a single spanning transmembrane protein that resides in apical protrusions, in both enterocytes [60] and tuft cells. Imaging revealed strong CDHR5 signal at the distal tips of protrusions as expected, and in large vesicles that were closely associated with Ac-tubulin containing microtubules in the sub-apical and perinuclear regions (Figure 7C). These results suggest that the cytoskeletal superstructure supporting the tuft could play a role in delivering or retrieving membranous cargoes to and/or from the apical surface.

**Figure 7.**
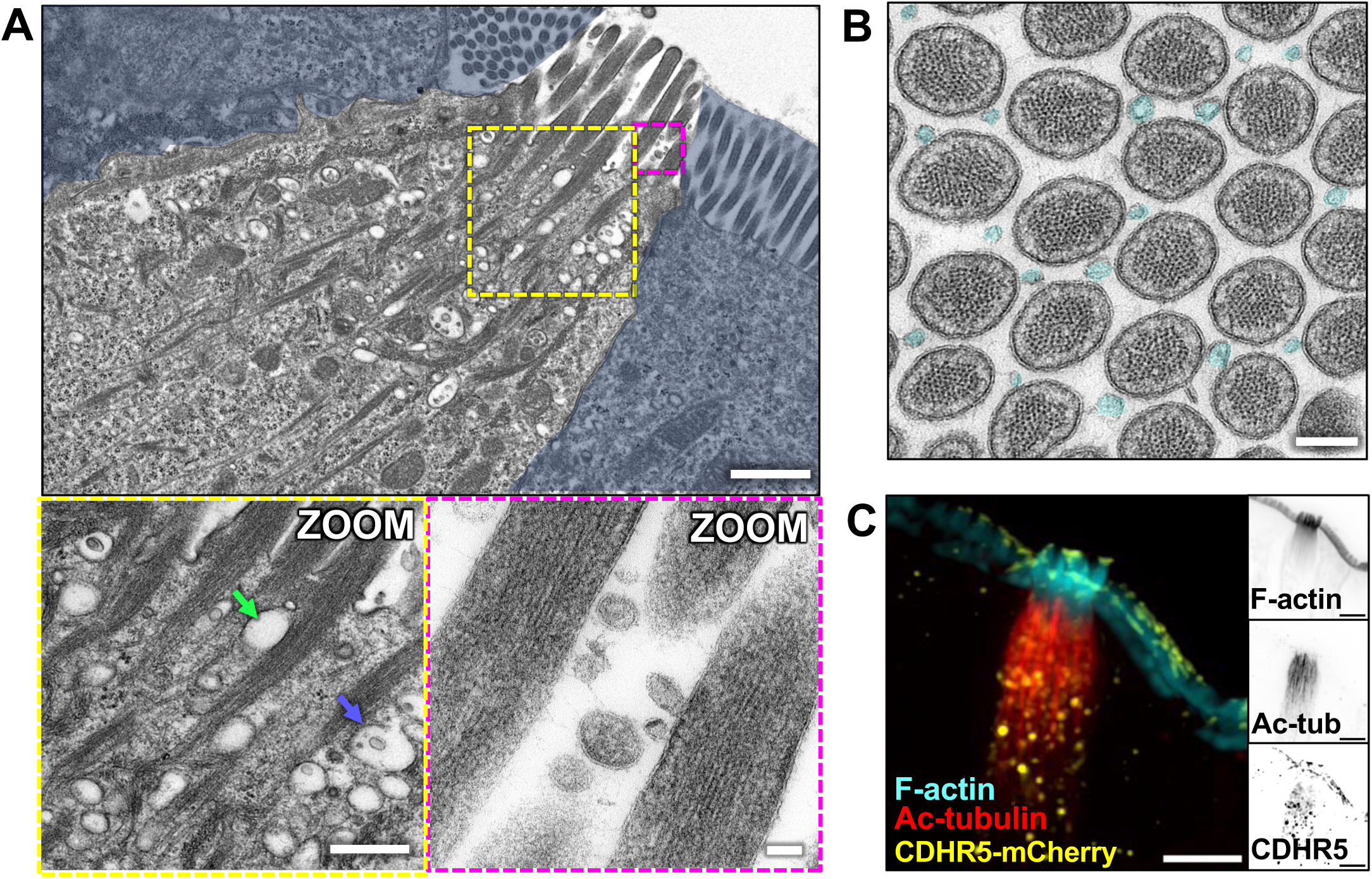
Membranous organelles are associated with the tuft cell cytoskeletal network. **A)** TEM of ultrathin tissue slice showing lateral tuft cell section (scalebar = 1 µm) with enterocytes masked in blue. Zoom inset (left) shows vesicles along core bundles near the apical surface (scalebar = 400 nm). Blue arrow points to a multivesicular body, green arrow points to electron lucent vesicle. Zoom inset (right) shows extracellular vesicles (EVs) between apical protrusions (scalebar = 50 nm). **B)** TEM of ultrathin tissue slice showing *en face* section of the apical tuft showing EVs (cyan) between protrusions (scalebar = 200 nm). This image was taken of the same tuft cell shown in Figure 2A. **C)** MaxIP SDC image of tuft cell from CDHR5-mCherry mouse, immunostained for mCherry to boost signal and acetylated tubulin (scalebar = 5 µm).

### Tuft cells have increased secretory and endocytic pathway markers relative to enterocytes

The dense collection of vesicles in the sub-apical cytoplasm and the presence of EVs outside the tuft suggest that tuft cells engage in robust membrane trafficking activities. We therefore stained tuft cells for markers of endo- and exocytosis. Remarkably, tuft cells exhibit robust enrichment of dynamin-2 beneath the apical surface (Figure 8A, B). Dynamin-2 drives vesicle scission at the plasma membrane during endocytosis and is also involved in the secretory pathway, promoting the release of vesicles from the Golgi network and the fusion of vesicles at the apical membrane [61]. We also stained for protein kinase C and casein kinase substrate in neurons 2 (pacsin-2), an F-BAR protein involved in apical endocytosis [62]; this factor was also markedly increased in tuft cells compared to neighboring enterocytes (Figure 8C, D). Together, these data suggest that tuft cells exhibit elevated trafficking activities and further imply that the tuft plays a fundamental role in transferring materials to and/or from the extracellular space.

**Figure 8.**
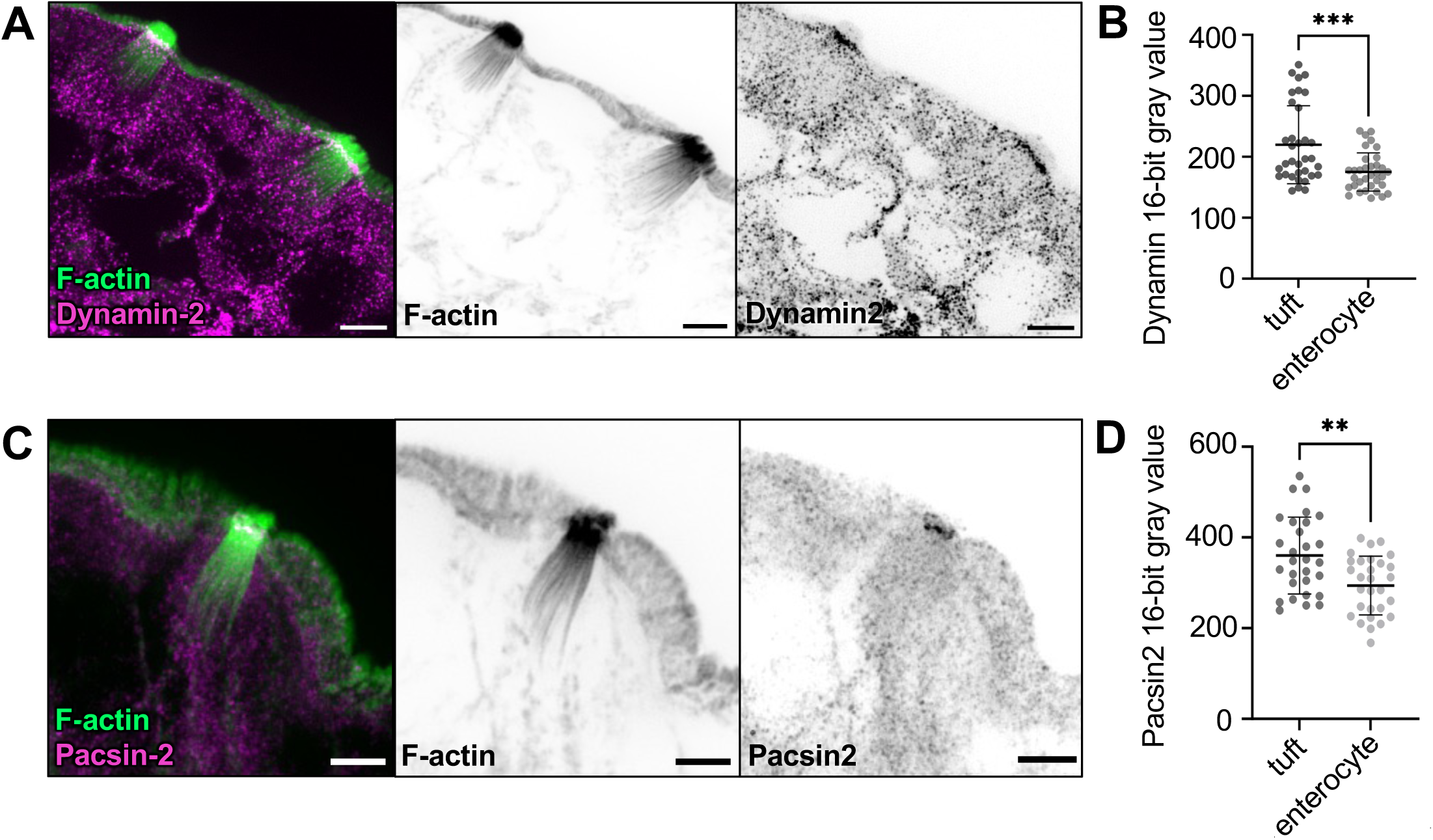
Tuft cells have increased secretory and endocytic pathway markers relative to enterocytes. **A & C)** MaxIP SDC image of lateral tissue section immunostained for dynamin-2 or pacsin-2 respectively. F-actin marked with phalloidin (scalebar = 5 µm). **B & D)** Mean intensity measurements taken from sum intensity projections of dynamin-2 (unpaired t-test, p = 0.0015, n = 28 tuft cells over 3 mice) and pacsin-2 (unpaired t-test, p = 0.0015, n = 34 tuft cells over 3 mice) in the apical surface of tuft cells versus enterocytes. Error bars denote mean ± SD.

## DISCUSSION

Whereas the physiological roles of canonical apical specializations such as the brush border and hair bundle are well established (solute uptake and mechanosensation, respectively) [63, 64], how the apical tuft supports the functions of the tuft cell remains unclear. We reasoned that by developing insight on the architecture of the tuft and its underlying cytoskeleton, we could begin to hypothesize how this structure might be leveraged *in vivo* to promote tuft cell function and intestinal homeostasis. Previous studies on tuft cells in diverse tissues offered glimpses of the cytoskeletal underpinnings of their unique morphology, although they stopped short of providing the specific molecular and structural details needed to understand the cell biological function of the tuft [2, 16, 17]. To fill these gaps, we combined light and electron microscopy to build detailed maps of the cytoskeletal structures and associated binding proteins that comprise the tuft. Below we discuss observations on the organization and composition of these structures, and their implications for understanding the subcellular function of the tuft.

### Organization of protrusions and giant core actin bundles in the tuft

Tuft protrusions extend from the apical surface in a cluster with a median packing angle of ∼59° when viewed *en face*, indicative of tight, hexagonal packing. For cylindrical objects, a hexagonal arrangement allows the cell to achieve maximum packing density. However, the spread of packing angles observed throughout the tuft was large (range of 20°-120°) revealing flexibility in how giant actin bundles are situated relative to their neighbors. How giant actin bundles are positioned and organized during tuft cell differentiation remains unclear. Enterocyte microvilli exhibit near perfect hexagonal packing, which is enforced by protocadherin-based intermicrovillar adhesion complexes (IMACs) that localize to the distal tips of these protrusions [60]. Interestingly, some IMAC components, such as CDHR5, are expressed in tuft cells and positioned to serve a similar role (Figure 7C). Whether tuft cells express and localize a full complement of IMAC proteins to control the packing of tuft cell protrusions remains an open question for future studies. However, our dataset does suggest additional mechanisms that might control the organization of tuft protrusions. For example, tight packing could be the result of compressive forces applied by the junctional contractile ring of F-actin and NM2, which encircles the cluster of giant core actin bundles (Figure 1N, O). This general idea is supported by our quantitative morphometry, which showed that the cross-sectional areas of the cell and the tuft converge right at the transition point where giant actin bundles emerge as protrusions (Figure 1H). Measuring such compressive forces directly is not possible, but high levels of contractility in this belt can be implied from the smaller *en face* areas and circularity of the apical perimeter. In enterocytes, NM2A and NM2C localize to a similar circumferential belt, but are also found in a medial plane that extends throughout the terminal web [31, 65], a meshwork of actin and intermediate filaments found just below the apical surface. Although previous EM studies were unable to resolve a terminal web meshwork in tuft cells [16], we were able to detect both NM2 variants in this plane. This sub-apical population of myosins might also constrain the packing of giant core actin bundles, either through direct binding/crosslinking or by propagating constraining mechanical forces generated by the circumferential belt across the sub-apical region.

### Filament organization in giant core actin bundles

Previous studies highlighted EPS8 as a barbed-end binding protein that is enriched in discrete puncta at the distal tips of actin-bundle supported protrusions including microvilli, stereocilia and filopodia [38, 40, 66]. Although we only detected low levels of EPS8 at the distal tips of tuft protrusions, a structurally related family member, EPS8L2, strongly localized to the tips (Figure 3), suggesting that filaments within giant core bundles are uniformly oriented (i.e. polarized) in a barbed-end out configuration. The strong distal end enrichment of EPS8L2 is also consistent with a model where individual filaments extend continuously through the full length of the core bundle, although TEM confirmation of this point was complicated by the tilt and curvature of these structures throughout the tuft, which made it difficult to capture their full length in ultra-thin sections.

TEM imaging of giant core bundle cross-sections revealed that these structures are composed of ∼100 hexagonally packed actin filaments. This highly ordered pattern contrasts with the F-actin core found in hair cell stereocilia, which exhibit poorly ordered ‘fluid’ packing, characterized by tight but irregular spacing of filaments [67]. The hexagonal packing observed in tuft cell giant core bundle might reflect the kinetics or mechanism of bundle polymerization, as highly ordered structures are generally produced through slow growth and low actin bundler concentrations [68]. The details of filament packing are important as they may ultimately dictate the mechanical flexibility of the bundle in response to lateral forces [69]. In stereocilia, filaments are crosslinked with multiple structurally distinct actin bundlers (e.g. fascin-2, plastin-1, espin-1) and fluid packing is the result of filaments accommodating these different crosslinker lengths [67]. Indeed, knockout of the most abundant crosslinker in hair cells, plastin-1, shifted filament organization from fluid to well-ordered hexagonal packing [67]. Based on those results, the ordered packing of filaments in giant core actin bundles observed here implies that the activity of single actin crosslinker dominates throughout these structures.

### Regionalization of actin binding proteins along the giant core bundle axis

Although our ultrastructural evidence argues in favor of a single crosslinker in giant core actin bundles, our immunofluorescence imaging detected several bundlers in these structures, including advillin, espin, and LIMA1 (Figure 4). These findings add to the short list of actin binding proteins, including fimbrin and girdin, which have been localized to the tuft by others [2, 21]. Our analysis does not allow us to rank order the abundance of the factors we examined. However, these proteins did exhibit an unexpected regionalization along the giant bundle axis, with advillin and espin occupying the membrane-wrapped distal ends, and LIMA1 demonstrating restricted localization to the more proximal, rootlet ends. Thus, filament packing patterns could be dictated locally by the specific complement of bundlers present in that segment. Such regionalization would also allow the tuft cell to create segment-specific mechanical properties, which may be important for function. The rootlet localization of LIMA1 is reminiscent of MISP, an actin bundling protein that exhibits specific enrichment on the rootlets of enterocyte microvilli [50]. Recent studies showed that loss of MISP function in an epithelial cell culture model shortened rootlets, whereas MISP overexpression led to rootlet hyper-elongation, through a mechanism involving competition with membrane-actin linker Ezrin [50]. Interestingly, previous *in vitro* studies showed that LIMA1 holds actin cross-linking activity, slows F-actin depolymerization, and inhibits binding of the Arp2/3 complex responsible for branched filament nucleation [70]. All of these activities are consistent with a role in rootlet elongation. Mechanistic follow-up studies will be needed to define the function of LIMA1 in shaping these unique structures.

### A cytoskeletal superstructure supports the tuft

By generating high-resolution 3D image segmentations, we discovered that microtubules and giant core actin bundles interdigitate and exhibit striking co-alignment throughout the sub-apical cytoplasm, with regions of overlap stretching for several microns. Such co-alignment is highly suggestive of specific molecular cross-linking between the two cytoskeletal networks, which could further constrain bundle spreading in the sub-apical compartment (Figure 1G, H). Physical contact between microtubules and actin networks is known to be critical for normal cell function in a range of tissue contexts [71]. Neuronal growth cones offer a striking example; in this system, intimate cross-linking between the two cytoskeletons is essential for neurite outgrowth, growth cone motility, and steering [72]. However, in that case, a central bundle of microtubules connects with a peripheral meshwork of F-actin that is highly branched and extends to the leading edge to drive cell motility. The interdigitated microtubules/actin bundle super-structure revealed here is morphologically unique and likely holds important implications for understanding the physiological function of tuft cells (more below).

### Potential functions for the cytoskeletal superstructure

The giant core actin bundles that comprise the tuft extend their rootlets many microns through the sub-apical cytoplasm. Such cytoskeletal architecture is rare in animal cell biology, although it does point to exciting possibilities for potential functions. For example, in any cytoskeletal network, the net orientation of polymers dictates the direction of motor protein driven transport [73]; polarized (i.e. uniform) filament orientation allows for unidirectional movement of motors, whereas mixed filament polarity can lead to motor stalling as demonstrated *in vitro* for MYO10 [74]. From this perspective, the parallel and polarized barbed-end out orientation of actin filaments in giant core bundles seem well suited for supporting efficient myosin-driven transport. Although MyTH4-FERM domain containing myosins are believed to drive transport within the confines of surface protrusions such as filopodia (MYO10) [36], microvilli (MYO7B)[75, 76], and stereocilia (MYO7A, MYO15A) [75, 76], we are not aware of any precedent in the literature for myosin-dependent transport along large parallel actin bundles in open cytoplasm. Investigating this idea in tuft cells would first require identification of candidate myosin motors holding structural and kinetic properties that are compatible with a role in transport (e.g. a dimeric structure with a processive mechanochemical cycle) [77]. Another possibility is that the cytoskeletal superstructure supports directed transport by microtubule motors (i.e. kinesin family members or dynein), which have well established roles in cytoplasmic trafficking [78]. In this scenario, the long rootlets of giant core bundles might simply serve as a physical scaffold for the alignment of stable microtubules. As some myosins directly interact with kinesins and such interactions increase the movement of both motors along networks containing F-actin and microtubules [79], cooperation between these systems might also be possible in the cytoskeletal superstructure that supports the tuft.

What cargoes might be transported along the tuft cytoskeleton? Tuft cells are responsible for the secretion of several effectors including IL-25, CysLTs, PGD2, and Ach, [9–12] which could be potential cargoes. EM and light microscopy analysis also revealed large numbers of vesicles with diverse morphologies distributed along giant core bundle rootlets. In some cases, these were identifiable as MVBs, leading to the possibility that these organelles are trafficked apically, along rootlets, to eventually release of their vesicle cargoes from the cell surface. The abundance of EVs we observed between individual tuft protrusions is consistent with this idea. Live imaging of tuft cells expressing labeled versions of secreted effectors or MVB markers will be needed to determine if the cytoskeletal superstructure in the tuft supports these trafficking functions.

### Next steps

Although previous studies uncovered general characteristics of tuft cells, the quantitative morphometry we report here provides a framework for future mechanistic studies of tuft cell function. Additionally, our work provides an experimental blueprint and quantitative point of comparison for eventual characterization of tuft cells in other tissues, as well as neuron-like and immune-like tuft cell subtypes. Tuft cells have been increasingly implicated in intestinal health for parasite clearance as well as restoration of intestinal barrier function in models of Crohn’s disease and ulcerative colitis [26]. Thus, investigation of how the unique tuft cell cytoskeletal features identified here are perturbed in these disease states should be an important goal for future studies.

## METHODS

All authors had access to the study data and reviewed and approved the final manuscript.

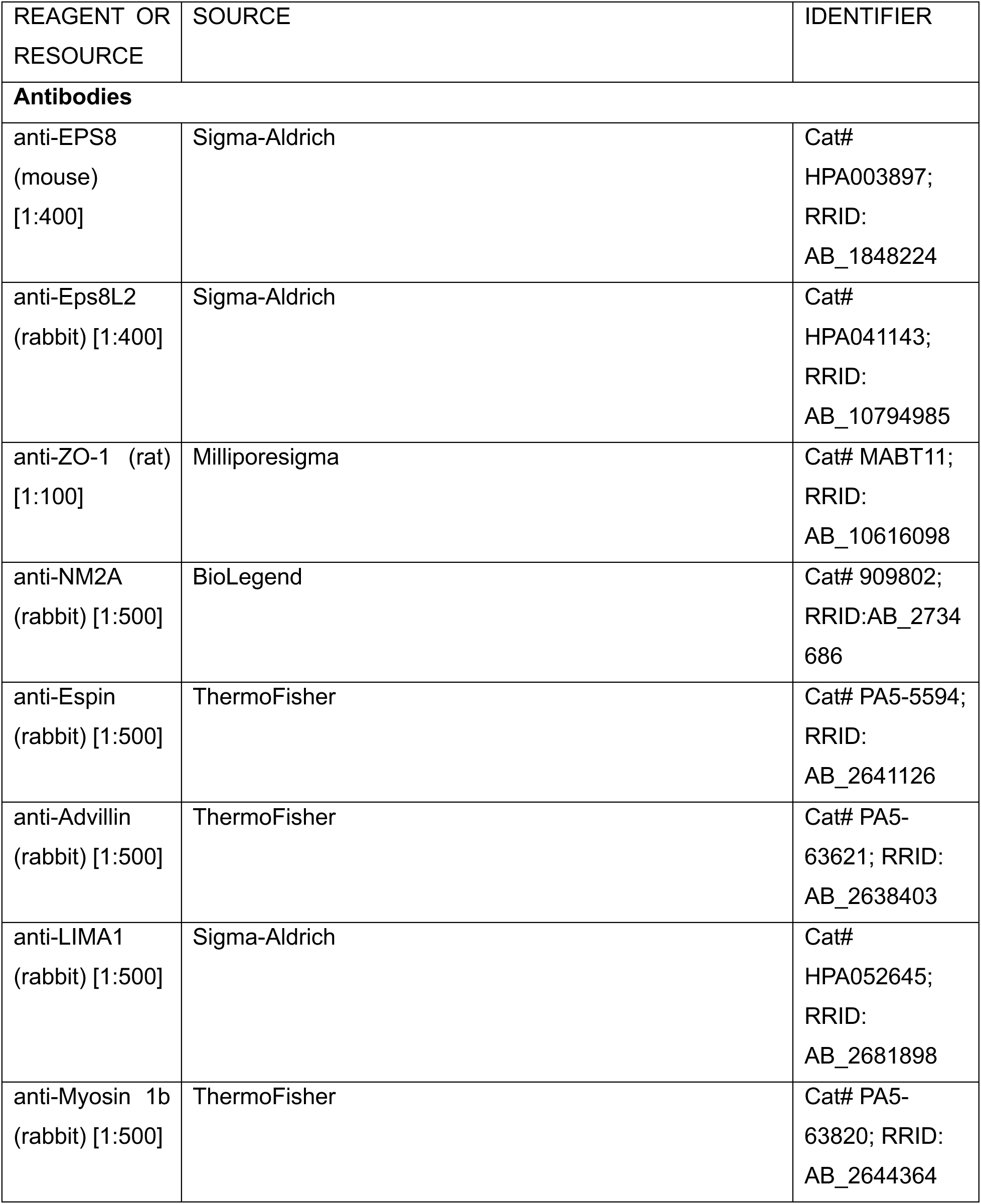
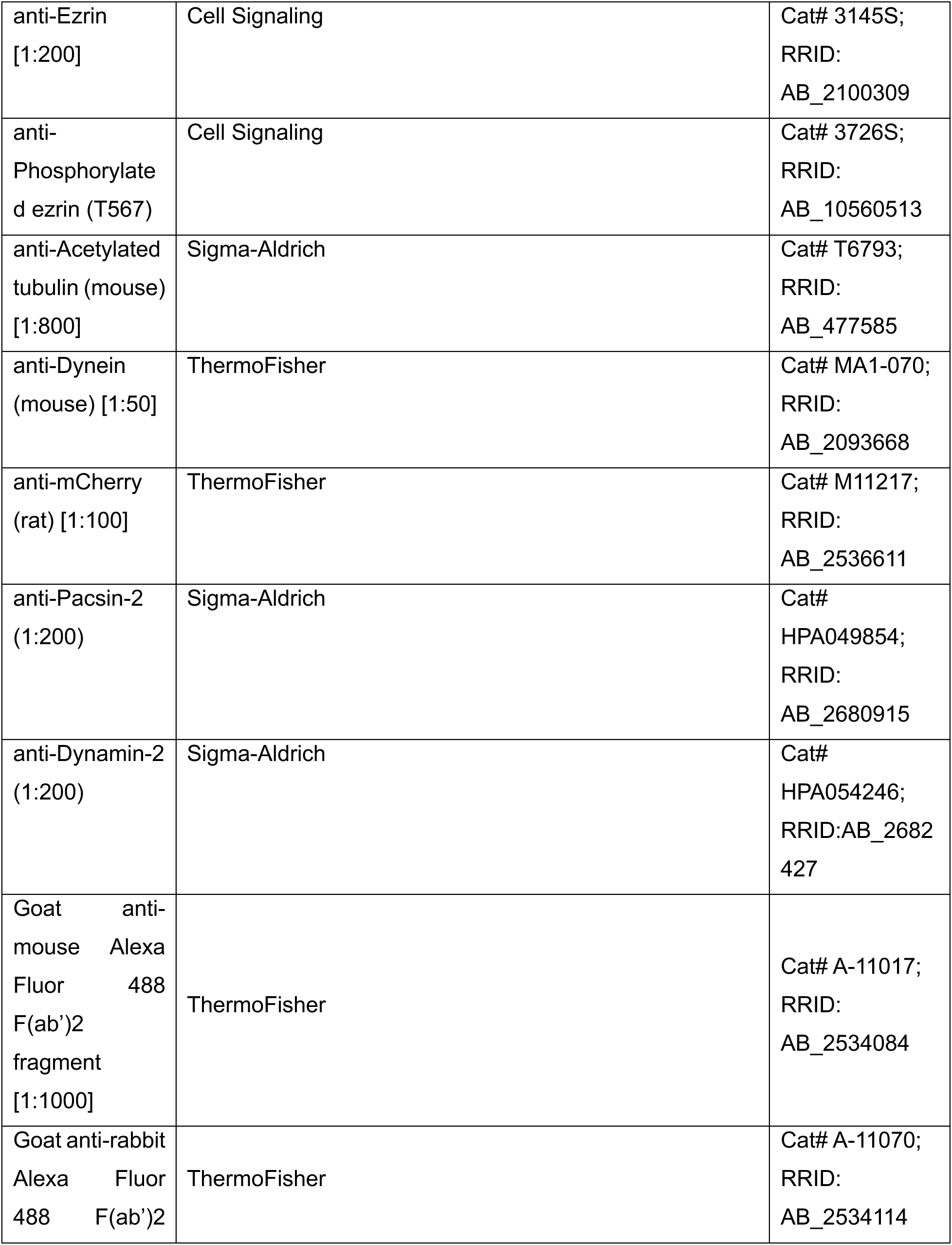
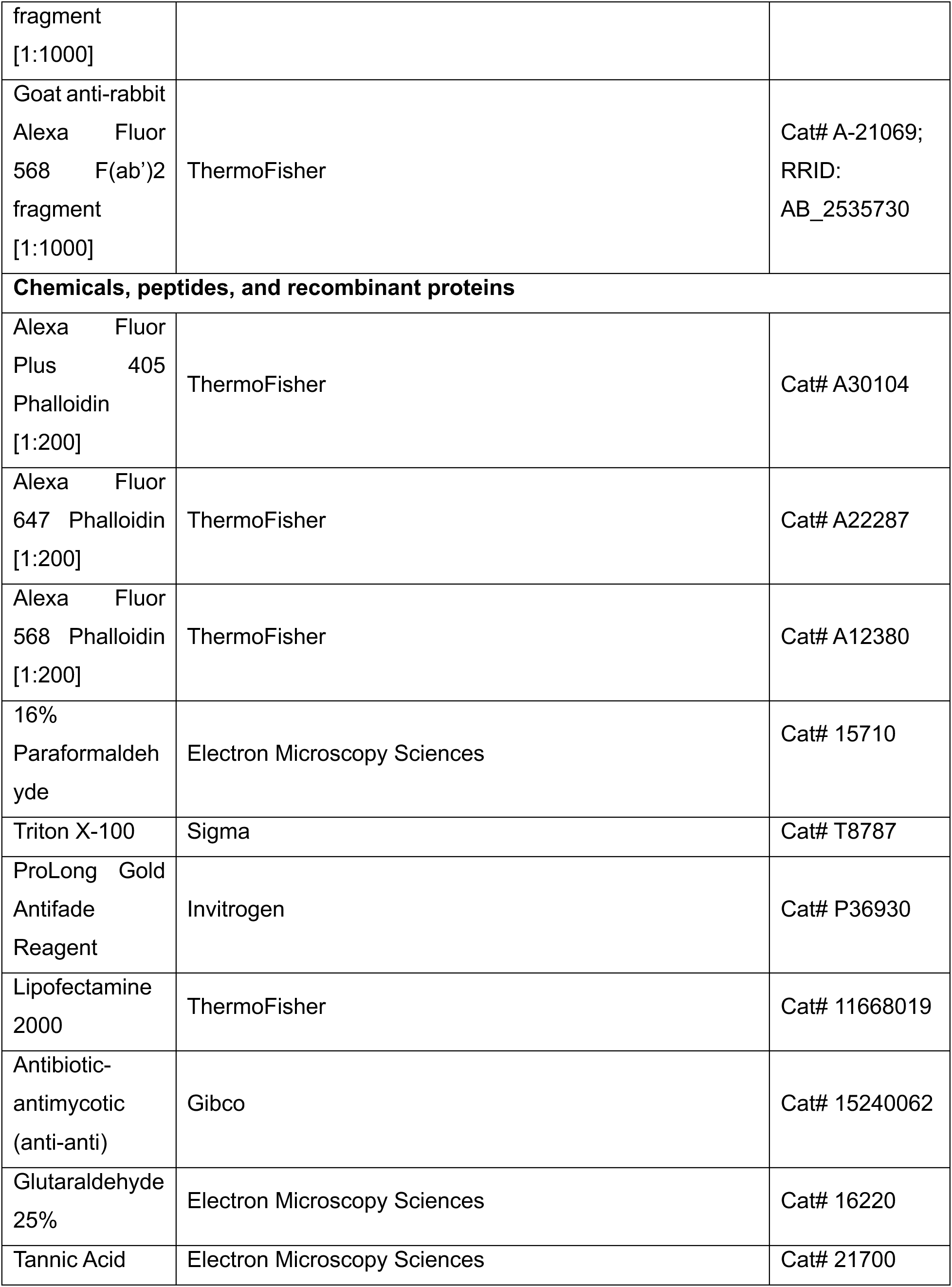
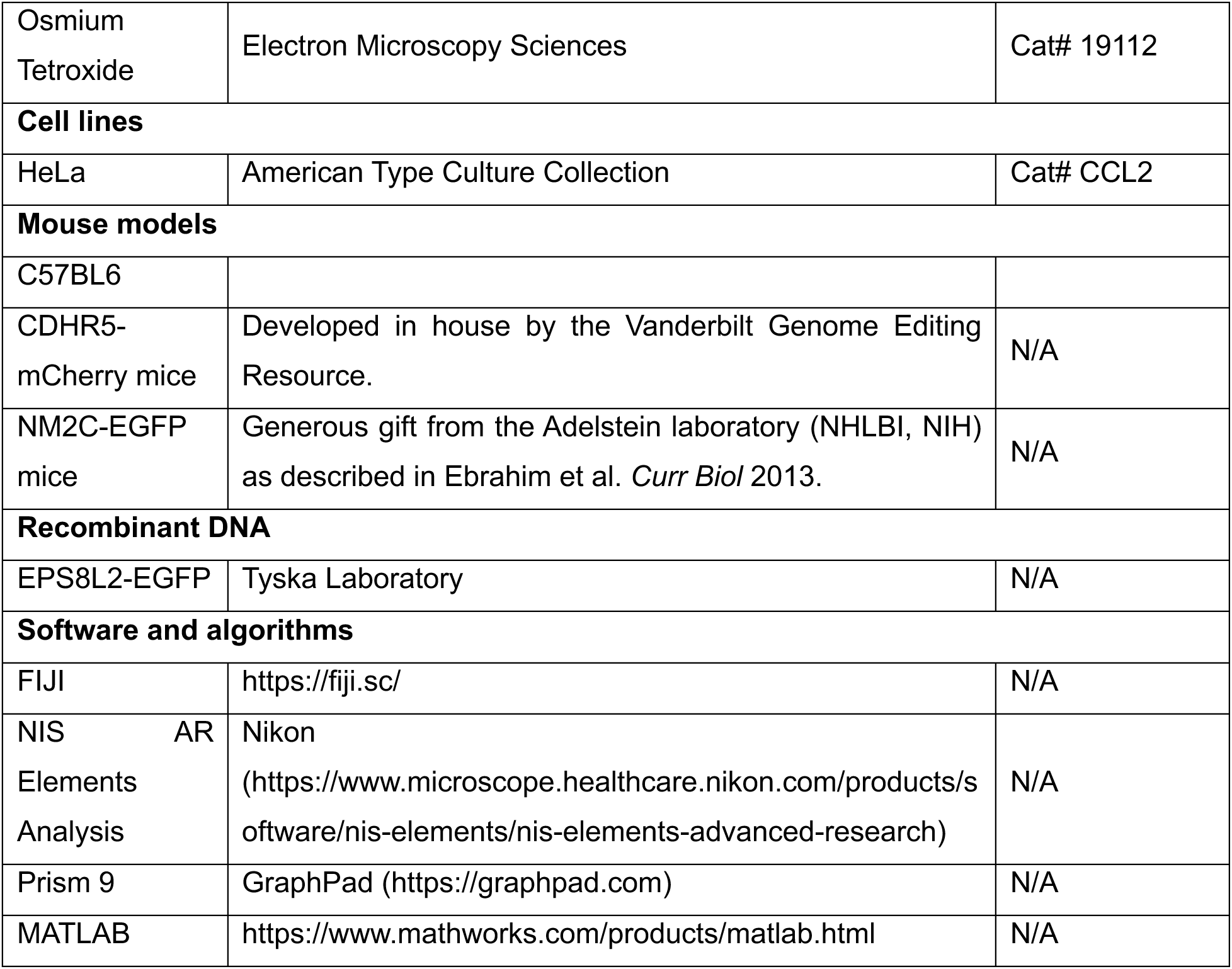

### EXPERIMENTAL MODEL DETAILS

#### Cell culture model

HeLa cells were cultured at 37°C and 5% CO2 in Dulbecco’s modified Eagle’s medium (DMEM) (Corning #10-013-CV) with high glucose and 2 mM L-Glutamine supplemented with 10% fetal bovine serum (FBS). 20 Transfections were performed using Lipofectamine 2000 (Thermo Fischer #11668019) 21 according to the manufacturer’s protocol. Cells were replated onto 35mm glass-bottom dishes or coverslips (Cellvis #D35-20-1.5-23 N) and incubated in Lipfectamine overnight. Media was replaced the following morning and cells were washed gently in PBS.

#### Mouse models

Animal experiments were carried out in accordance with Vanderbilt University Medical Center Institutional Animal Care and Use Committee guidelines under IACUC Protocols. NM2C-EGFP mouse were obtained as a generous gift from Dr. Robert Adelstein (NHLBI, NIH) and were described previously [31]. Mice expressing endogenously tagged CDHR5 mice were created in collaboration with the Vanderbilt Genome Editing Resource. Briefly, CRISPR/Cas9 genome engineering methods were used to insert a flexible linker and mCherry coding sequence at the 3’ end of the CDHR5 terminal coding exon.

### METHOD DETAILS

#### Succinate administration

For tissue used in TEM preparation, 120 nM of sodium succinate hexahydrate (with dextrose at 1%/volume to improve taste) was added to drinking water of wild-type (C57BL6) mice for one week prior to euthanasia and tissue preparation.

#### Frozen and whole mount tissue preparation

Segments of proximal intestine were removed, flushed with phosphate-buffered saline (PBS), and prefixed for 15 min with 4% paraformaldehyde (PFA) to preserve tissue structure. The tube was then cut along its length, subdissected into 0.5 mm–chunks, fixed for an additional 30 min in 4% PFA at room temperature, and washed three times in PBS. Whole mount samples were then moved to Eppendorf tubes for staining. For frozen sections, tissue samples were gently placed on top of a 30% sucrose solution in TBS and allowed to sink to the bottom overnight at 4°C. Specimens were then swirled in three separate blocks of OCT (Electron Microscopy Sciences), oriented in a block filled with fresh OCT, and snap frozen in dry ice-cooled acetone. Samples were cut into 10-μm sections and mounted on slides for staining.

#### Immunofluorescence

##### Cell culture

For spinning disk confocal imaging, cells were washed three times with pre-warmed phosphate-buffered saline (PBS) before being fixed in 4% paraformaldehyde (Electron Microscopy Sciences #15710) in PBS for 15 mins at 37°C. Cells were then incubated for 1 hr with Alexa Fluor 568 phalloidin (1:200; Invitrogen #A12380) at room temperature. Coverslips were then washed three times in PBS and mounted on slides with 20 μL ProLong Gold (Invitrogen #P36930).

##### Frozen tissue sections

For Zeiss Airyscan or spinning disk confocal imaging, frozen tissue sections were rinsed three times in PBS and permeabilized with 0.1% Triton X-100 in PBS for 15 mins at room temperature. After permeabilization, tissue slides were rinsed three times in PBS and blocked with 10% BSA (Research Products International #9048-46-8) in PBS for 2 hr at 37°C in a humidified chamber. Immunostaining was performed using primary antibodies (see table for details) diluted in 1% BSA overnight at 4°C. After incubation with primary antibody, slides were rinsed three times in PBS and incubated for 2 hr with appropriate secondary antibodies (see table for details) and an Alexa Flour conjugated phalloidin (see table for details) at room temperature. Slides were then washed three times with PBS and coverslips were mounted with 60 μL ProLong Gold (Invitrogen #P36930) overnight.

##### Wholemount tissue sections

For spinning disc confocal imaging, whole mount tissue was stained in Eppendorf tubes. Tissue was rinsed three times in PBS and permeabilized with 0.2% Triton X-100 in PBS for 30 minutes at room temperature on a rocker. After permeabilization, tissue was rinsed three times in PBS and blocked in 5% BSA overnight at 4°C on a rocker. Primary antibody (see table for details) diluted in 1% BSA was added and left overnight at 4°C on a rocker. The next day, the tissue was washed three times in PBS and the appropriate secondary antibodies (see table for details) were diluted in 1% BSA for 4 hours at room temperature on a rocker. At two hours of incubation, an Alexa Fluor conjugated phalloidin was added to the tissue (see table for details). The tissue was then washed three times in PBS and mounted villi-down on a glass-bottom dish with 20 μL ProLong Gold with a coverslip on top overnight.

#### Light microscopy

Laser scanning confocal microscopy was performed on a Nikon A1 microscope equipped with 488 nm, 568 nm, and 647 nm LASERs and a 100x/1.49 NA TIRF oil immersion objective. Spinning disk confocal imaging was conducted using a using a Nikon Ti2 inverted light microscope with a Yokogawa CSU-W1 spinning disk head, a Photometrics Prime95B sCMOS camera, four excitation LASERs (488, 568, 647, and 405 nm), and a 100X/1.49 NA TIRF oil immersion objective or a 60X/1.49 NA TIRF oil immersion objective. Images presented in figures were deconvolved (Richardson-Lucy deconvolution of image volumes, 20 iterations) using Nikon Elements software. Super-resolution images were collected on a Zeiss LSM980 Airyscan microscope with four excitation LASERs (488, 568, 647, and 405 nm,),and a 63X/ 1.43 NA oil immersion objective; images were processed using Zeiss Zen software.

#### Electron microscopy

To prepare samples for TEM, 1 mm pieces of small intestine were fixed for 1 h in 2% paraformaldehyde, 2% glutaraldehyde, in 0.1 M cacodylate buffer with 2 mM CaCl_2_. The samples were washed in 0.1 M HEPES and slowly equilibrated with 30% glycerol as a cryoprotectant. Samples were plunge frozen in liquid ethane followed by freeze-substitution in 1.5% uranyl acetate in methanol for 48 hours at −80C. Samples were washed extensively in methanol and infiltrated with HM20 Lowicryl. The HM20 was polymerized by UV light at −30C for 24 hours under nitrogen vapor. Samples were sectioned on a Leica UC7 ultramicrotome with a nominal thickness of 70 nm on 200 mesh Ni grids and poststained with uranyl acetate and lead citrate. Images were collected with a Tecnai T-12 transmission electron microscope operating at 100 kV using an AMT nanosprint5 CMOS camera.

#### Analysis of scRNAseq data

We used a previously published scRNAseq dataset to probe for tuft cell enriched candidate genes [26]. This dataset was generated using droplet-based single cell sequencing from the small intestine of 6 wild-type mice. Further method details can be found in the original paper. UMAPs were displayed using CellxGene, a single-cell visualization platform.

#### Image analysis and statistics

All images were processed using Nikon Elements or FIJI software (https://fiji.sc/).

##### Actin bundle length and intensity, and localization of actin-binding proteins

Lateral images of frozen sections were analyzed using FIJI. Maximum intensity projections of core actin bundles were generated and lines were drawn from the apical tuft to the bottom of the rootlets to enable measurements of bundle length and marker intensity along the bundle axis.

##### Number, angle, and bundle spreading measurements

Images captured *en face* using whole mount tissue were used for these quantifications. Maximum intensity projections of image volumes acquired just beneath the apical surface were used to count the number of individual core actin bundles per tuft cell. The core actin bundles were identified by dark spot identification in Nikon Elements and angles were measured from every bundle to two adjacent neighbors. Bundle spreading was measured by calculating bundle area versus cell area (identified by phalloidin staining), using individual slices at the apical surface, 1.5 μm, 3 μm, and 4.5 μm below the apical surface.

##### Fourier analysis of TEM images

To map filaments of equidistant packing or hexagonal packing back onto an original TEM image, Fourier filtering was performed with DigitalMicrograph and Fiji. Target areas were selected and then filtered using an Fast Fourier Transform. A circular (for equidistant packing) or hexagonal mask (for hexagonal packing) was placed over the predominant reflections in Fourier space using the masking tools built-in DigitalMicrograph, making the mask as tight as possible to reduce background Fourier signal. After applying the mask, an inverse FFT was performed to generate the corresponding real-space image. This image was thresholded in Fiji to remove background signal, pseudocolored, and then merged with the original image.

##### Trainable WEKA segmentation of giant actin bundles and microtubules

Oversampled images of *en face* whole mount tissues stained with phalloidin and acetylated tubulin were opened in FIJI and subject to Trainable WEKA segmentation [29] to identify core actin bundles and microtubules independently. Actin probability maps were put under the minimum auto local thresholding while microtubule probability maps were put under the default auto local threshold. These thresholds were chosen to maximize selection of the cytoskeleton. Both maps were opened in Nikon Elements and thresholded so that individual objects (in this case individual core actin bundles or microtubules) could be visualized. From this data, we obtained the pitch (°) data. To determine the interaction between microtubules and core actin bundles, we dilated the core actin bundle signals and measured the length of overlap with microtubules. We further compared the volume of the overlap with the volume of the core actin bundles to determine the percent of actin polymer overlapping with microtubules.

##### Intensity analysis of antibody staining

For quantification of NM2C intensities in tuft cells and enterocytes, we used FIJI to analyze *en face* images from whole mount tissues. Sum intensity projections encompassing the apical surface were generated, an ROI was drawn to capture the tuft cells or a neighboring enterocyte, and the mean intensities were measured. For other intensity measurements, including NM2A, ezrin, phospho-ezrin, dynamin-2, pacsin-2, and dynein, lateral images of frozen tissue sections were analyzed in FIJI. Sum intensity projections were generated to include the full tuft, an ROI was drawn over apical protrusions, and the mean intensity of the marker was measured using an ROI tool. An identically sized ROI was used to measure the mean intensities from neighboring enterocytes.

##### Nearest neighbor measurements for filaments within individual core actin bundles

FIJI plugin TrackMate was used to mark individual actin filaments within single giant core bundles. The center of each filament was exported as an x,y coordinate, and a custom MATLAB script (created with generative AI) was used to identify: (i) the number of nearest neighboring filaments within a 12 nm radius of each filament, (ii) the distance between nearest neighbors, and (iii) the angle between a filament and two adjacent nearest neighbors.

---

**Supplemental Figure S1.**
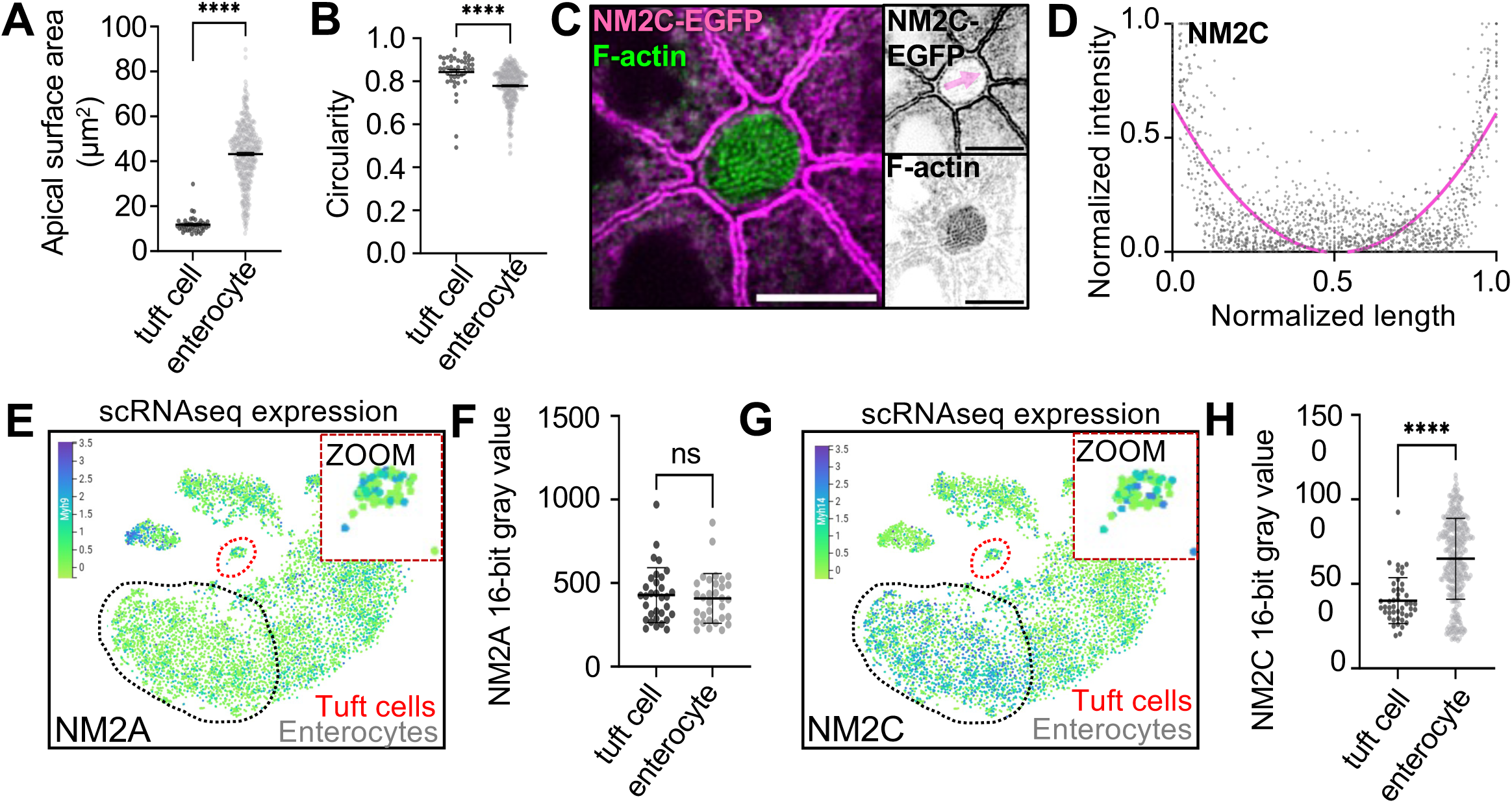
**A)** Apical surface area measurement using ZO-1 staining in whole mount tissue as imaged in Figure 1M. Unpaired t-test, p <0.001 (n = 45 tuft cells over 3 mice) **B)** Apical circularity measurement using ZO-1 staining in whole mount tissue as imaged in Figure 1M, unpaired t-test, p <0.001. **C)** MaxIP SDC image of *en face* whole mount NM2C-EGFP tissue, actin stained with phalloidin (scale bar = 5 µm). **D)** Graph of linescans for NM2C intensity drawn across the apical surface of a tuft cell (magenta arrow in Figure S1B) in MaxIPs of NM2C-EGFP whole mount tissue (n = 43 tuft cells over 3 mice). **E)** UMAP displaying ScRNAseq data of mouse intestinal tissue showing NM2A intensity measurement in tuft cells red circle/zoom and enterocytes gray circle (n = 6 mice). **F)** Mean NM2A intensity measurements in tuft cells versus enterocytes, unpaired t-test, p = 0.6179, Error bars denote mean ± SD, (n = 32 tuft cells over 3 mice). **G)** UMAP displaying ScRNAseq data of mouse intestinal tissue showing NM2A intensity measurement in tuft cells red circle/zoom and enterocytes gray circle (n = 6 mice). **H)** Mean NM2C intensity measurement in tuft cells versus enterocytes, unpaired t-test, p <0.0001, Error bars denote mean ± SD, (n = 46 tuft cells over 3 mice).

**Supplemental Figure S2.**
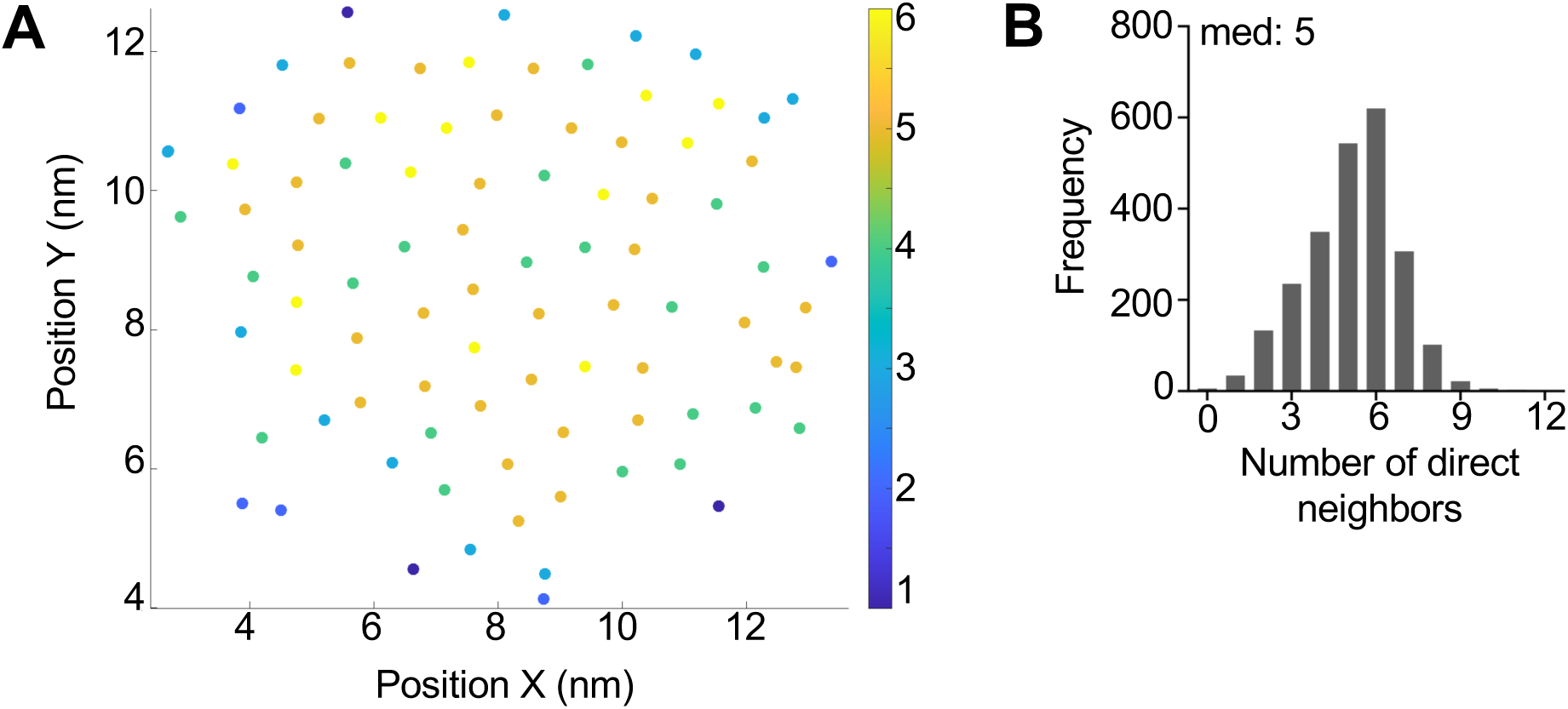
**A)** Example of nearest neighbors analysis showing a single giant core bundle and individual filaments. Filament coordinates were derived using Trackmate and put into a custom MATLAB script to quantify nearest neighbors (within a 12 nm radius). Filaments are color coded based on number of nearest neighbors. **B)** Frequency diagram of the number of nearest filament neighbors (within a 12 nm radius) (n = 22 bundles over 3 tuft cells).

**Supplementary Figure S3.**
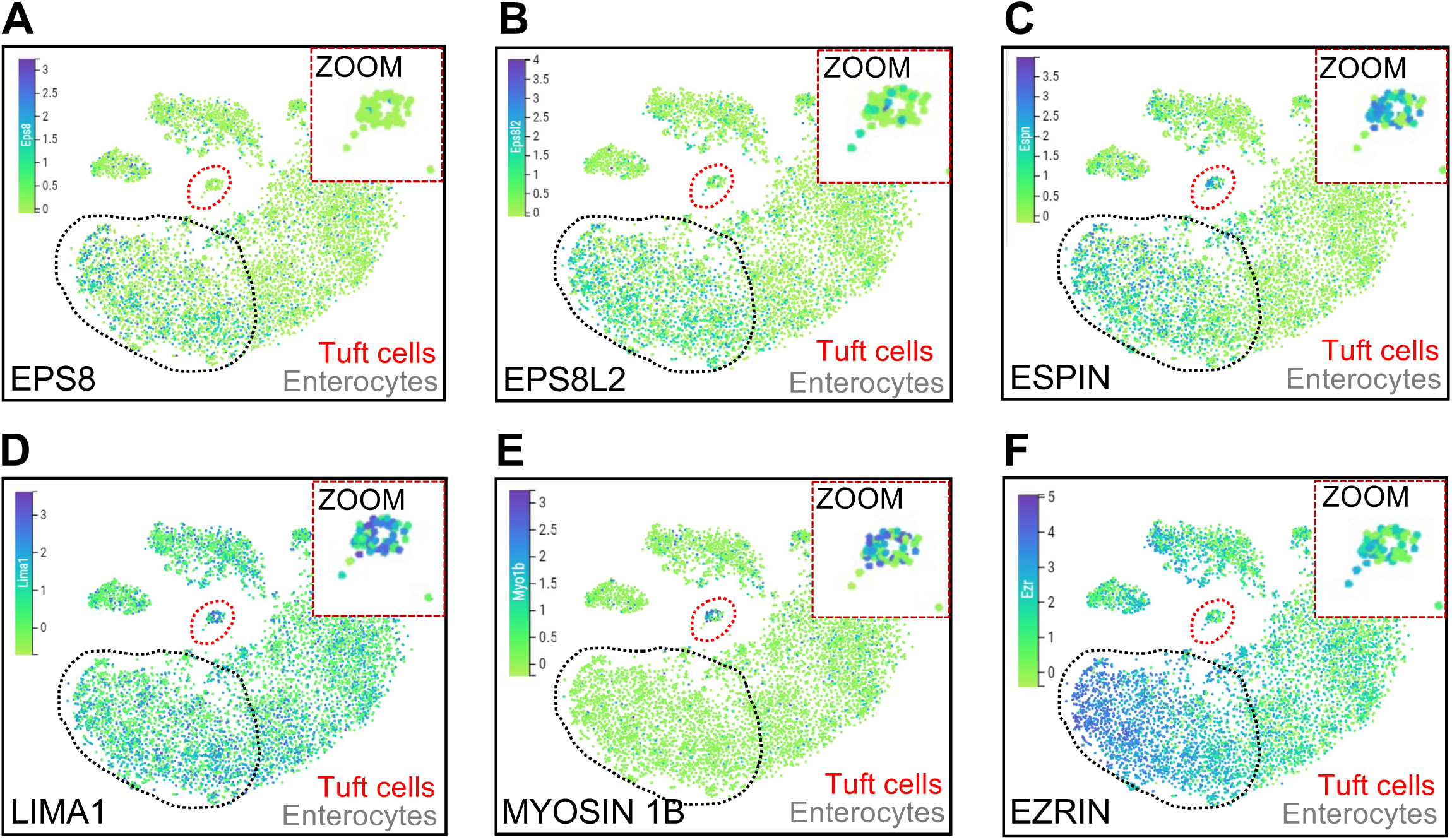
UMAPs showing ScRNAseq data of mouse intestinal tissue showing mRNA levels in **A)** EPS8, **B)** EPS8L2, **C)** espin, **D)** LIMA1, **E)** myosin 1b, & **F)** ezrin. Measurements in tuft cells red circle/zoom and enterocytes gray circle (n = 6 mice).

## Notes

**Grant support:** R01 DK103831 (KSL), R01 DK095811 (MJT), R01 DK111949 (MJT), R01 DK125546 (MJT)

### Competing Interest Statement

The authors have declared no competing interest.

